# USP37 protects mammalian cells during DNA replication stress by counteracting CUL2^LRR1^ and TRAIP

**DOI:** 10.1101/2024.09.03.610971

**Authors:** Fabrizio Villa, Johanna Ainsworth, Karim P.M. Labib

## Abstract

The USP37 deubiquitylase is important for mammalian cells to survive DNA replication stress but the underlying mechanisms are unknown. Here we demonstrate that USP37 binds the CDC45-MCM-GINS (CMG) helicase, which forms the stable core of the replisome and is regulated by ubiquitylation. The Pleckstrin-Homology Domain of USP37 binds CDC45 and structure-guided mutations that displace USP37 from CMG are sufficient to phenocopy loss of USP37 catalytic activity. Importantly, USP37 counteracts CMG helicase ubiquitylation by the CUL2^LRR1^ ligase, which induces helicase disassembly during termination. We show that depletion of CUL2^LRR1^ suppresses the sensitivity of Usp37 mutants to DNA synthesis defects and to ATR checkpoint kinase inhibitors. In contrast, mutation of the TRAIP ubiquitin ligase specifically suppresses the sensitivity of Usp37 mutants to topological stress during chromosome replication. We propose that USP37 evolved to reverse the untimely action of the two ubiquitin ligases that regulate the CMG helicase during chromosome replication in metazoa.

## Introduction

The eukaryotic replisome is built around the DNA helicase known as CMG (CDC45-MCM-GINS), which uses a hexameric ring of the MCM2-7 ATPases to encircle one of the two parental DNA strands at replication forks, in a highly stable fashion ^1,2^. The CMG helicase is assembled at origins of replication and must remain associated with each replication fork throughout elongation ^3^, since CMG is rapidly disassembled when displaced from DNA ^4,5^ and the helicase cannot be reassembled until the next cell cycle, to ensure that cells produce just one copy of each chromosome ^1,6^.

Disassembly of the CMG helicase is usually restricted to DNA replication termination ^7,8^. Recent work with budding yeast indicates that CMG helicase disassembly is an essential process ^9^, which nevertheless must be highly regulated to prevent it happening prematurely. The trigger for CMG disassembly is ubiquitylation ^10,11^, which leads to unfolding of the ubiquitylated Mcm7 subunit of CMG by the Cdc48 / p97 unfoldase and thus to disintegration of the helicase ^4,12,13^. Fungi and animal cells use different ubiquitin ligases to control CMG disassembly ^10,14,15^, the regulation of which has diversified considerably during the course of eukaryotic evolution.

Studies of DNA replication termination in *Caenorhabditis elegans*, *Xenopus laevis* egg extracts and mammalian cells indicated that CUL2^LRR1^ binds to the CMG helicase during DNA replication termination, triggering CMG ubiquitylation and disassembly ^14–17^. In contrast, the TRAIP ubiquitin ligase was shown in *Xenopus* egg extracts to associate with the replisome during elongation ^14^ and direct the ubiquitylation of proteins that are located in front of stalled replication forks ^18,19^, without causing CMG ubiquitylation *in cis*. In this way, TRAIP initiates the repair of Protein-DNA crosslinks ^18^, the resolution of transcription-replication conflicts ^20^, or the repair of inter-stand DNA crosslinks via *in trans* CMG ubiquitylation at a pair of converged replication forks ^19^, thereby providing access for DNA repair enzymes to the damaged DNA. In humans, mutations in TRAIP are associated with forms of microcephalic primordial dwarfism ^21^, which often reflect defects in regulation of the CMG helicase ^22–24^.

The activity of both CUL2^LRR1^ and TRAIP must be carefully regulated, to prevent the untimely ubiquitylation of the CMG helicase or of other targets. Until now, proteins that negatively regulate or counteract CUL2^LRR1^ and TRAIP had not been identified. In contrast, the DNA structure of replication forks was found to restrict CMG ubiquitylation by CUL2^LRR1^ to DNA replication termination ^4,5^. The DNA strand that is excluded from the CMG helicase at replication forks forms a steric block to CUL2^LRR1^ engagement with the helicase during elongation, until the moment when two converging CMG helicases from neighbouring origins have passed each other ^25^. However, it was unknown whether additional factors are required to counteract CUL2^LRR1^ at stalled DNA replication forks, where engagement of the helicase with the excluded DNA strand might be less robust, or might transiently be lost during DNA repair reactions. Moreover, the regulation of TRAIP activity is poorly understood and it was unclear whether additional mechanisms are needed to counteract TRAIP in response to defects in fork progression.

Here we show that binding of the USP37 deubiquitylase to the CMG helicase is a key determinant of survival when mammalian cells experience DNA replication stress, or are treated with inhibitors of the ATR checkpoint kinase. Our data identify USP37 as an important counterbalance to the CUL2^LRR1^ and TRAIP ubiquitin ligases in response to DNA synthesis defects and topological stress.

## Results

### The USP37 deubiquitylase associates with the CMG helicase in mammalian cells

To identify potential new regulators of the mammalian CMG helicase, we used CRISPR-modified mouse embryonic stem cells (ES cells) in which the SLD5 subunit of GINS was fused to the ‘Tandem Affinity Purification’ or TAP tag ^17^. The Protein A component of the tag binds to immunoglobulins, facilitating the specific and efficient isolation of TAP-tagged protein complexes from native cell extracts. Control mouse ES cells were grown in parallel with *TAP-Sld5* cells, either untreated or with a p97 inhibitor to prevent disassembly of ubiquitylated CMG during DNA replication termination ^17^. Extracts were incubated with antibody-coated magnetic beads, before mass spectrometry analysis of the material from four independent experiments (an example of the isolated material is shown in Figure S1A). The data confirmed the presence of all previously characterised subunits and partners of the CMG helicase (Tables S1-S4), together with additional factors such as PARP1, PIF1 (Figure S1B) and RTEL1 that function as accessory DNA helicases at replication forks, consistent with recent findings ^26–30^.

Two regulators of ubiquitylation were found to co-purify with CMG (Tables S1-S4). The first was the TRAIP ubiquitin ligase that we and others had previously found to interact with the CMG helicase in mammalian cells ^17,21^. The second was the deubiquitylase USP37, which was not known to associate with CMG but is present on replicating chromatin in *Xenopus* egg extracts ^14^ and mammalian cells ^31^, suggesting a potential role at DNA replication forks. Consistent with this view, depletion of USP37 in human cells by siRNA or CRISPR-Cas9 leads to increased stalling of DNA replication forks ^32–34^, genome instability ^35–38^, and sensitivity to agents that induce DNA replication stress ^32,34,39^. These studies led to the view that USP37 is an important regulator of DNA replication in mammalian cells but the associated molecular mechanisms remained unclear.

Immunoblotting confirmed the co-purification of USP37 with the CMG helicase (Figure S1C). To study the association of USP37 with CMG in more detail, we first used CRISPR-Cas9 to introduce the GFP tag into the *Usp37* locus in mouse ES cells (Figure S2A-C and E). The association of GFP-USP37 with the CMG helicase was not dependent upon ubiquitylation of the MCM7 subunit of CMG (Figure 1A, compare lanes 6 and 8, -/+ p97i), nor did it require the catalytic activity of USP37 (Figure 1A, lane 7; Propargylated ubiquitin or Ubi-Prg is a covalent inhibitor of cysteine deubiquitylases such as USP37 ^40^). These findings suggested that USP37 binds to the CMG helicase or an associated factor, rather than simply associating with polyubiquitin chains on CMG-MCM7.

**Figure 1.**
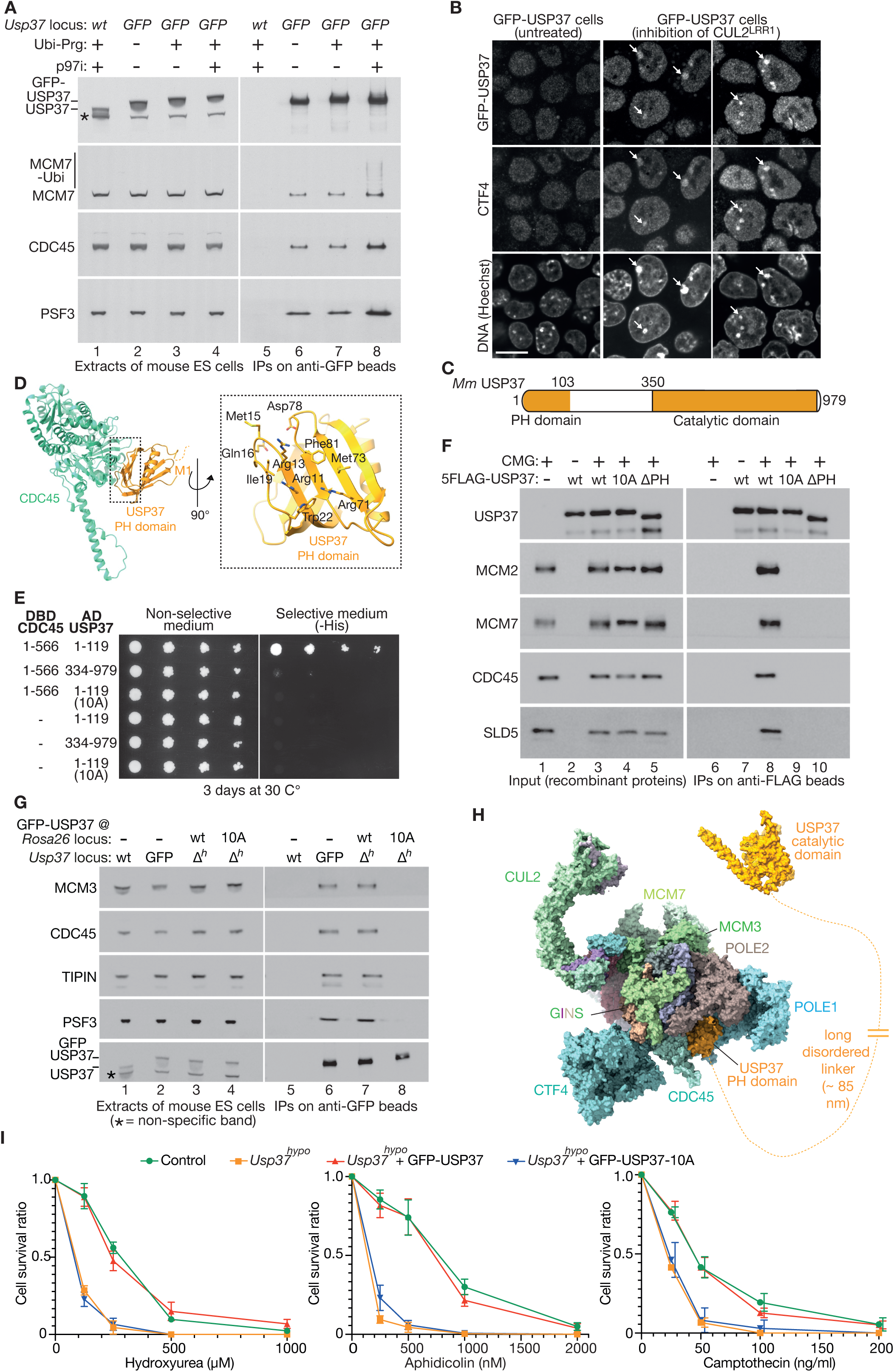
The Pleckstrin-Homology domain of USP37 couples the deubiquitylase to the CMG helicase and protects cells during DNA replication stress. (**A**) Mouse ES cells of the indicated genotype were treated with 5 µM CB-5083 (p97 inhibitor = p97i) as shown. Cell extracts were prepared in the presence of 5 μM Propargylated-Ubiquitin (Ubi-Prg = deubiquitylase inhibitor) as indicated, before incubation with anti-GFP beads. Bound proteins were monitored by immunoblotting. (**B**) GFP-Usp37 mouse ES cells were treated as indicated to inhibit CUL2^LRR1^ (see Materials and Methods for details), before analysis of fixed cells by spinning disk confocal microscopy. The arrows indicate the accumulation of CTF4 and GFP-USP37 on patches of constitutive heterochromatin, in response to inhibition of CUL2^LRR1^. Scale bar = 10 µm. (**C**) Outline of mouse USP37 protein. (**D**) AlphaFold Multimer prediction of the interaction between mouse CDC45 and the Pleckstrin-Homology (PH) domain of USP37. The panel on the right shows USP37 residues that are predicted to contact CDC45 and that were mutated in the USP37-10A allele. (**E**) Yeast two hybrid analysis of the interaction between mouse CDC45 and the USP37-PH domain. Budding yeast cells were transformed with plasmids expressing the indicated fusion proteins (DBD = Gal4 DNA Binding Domain, AD = Gal4 Activation Domain). Serial dilutions of transformed cells were spotted on the indicated medium, before growth for three days at 30°C. A positive two-hybrid interaction brings together the DBD and AD domains of Gal4 and leads to expression of the yeast *HIS3* gene, which allows growth on medium lacking histidine (-His). (**F**) Recombinant mouse CMG helicase and the indicated versions of FLAG-tagged USP37 were mixed in vitro, before incubation with magnetic beads coated with anti-FLAG antibody. The indicated factors were monitored by immunoblotting. (**G**) Extracts of mouse ES cells, with the indicated versions of the endogenous *Usp37* locus (wt = wild type, GFP = *GFP-Usp37*, Δ*^h^* = *Usp37^hypo^*) and with GFP-USP37 or GFP-USP37-10A expressed from the *Rosa26* locus as shown, were incubated with anti-GFP beads. The bound proteins were monitored by immunoblotting. (**H**) Model illustrating how the USP37-PH domain tethers the USP37 catalytic domain to the replisome via a long disordered linker. The interaction of the USP37-PH with CDC45 was modelled by AlphaFold Multimer and the resulting prediction was docked onto the cryo electron microscopy structure of the human replisome bound to CUL2^LRR1^ (PDB complex 7PLO). Length of the linker (residues 104-349 of USP37) was estimated using a contour length of 3.6 Angstroms per amino acid ^62^. (**I**) Mouse ES cells of the indicated genotypes were grown in the presence of hydroxyrea, aphidicolin or campothecin as shown for 24 hours, before growth for a further seven days in medium lacking hydroxyurea. Colonies were stained with crystal violet and the data were quantified as described in STAR methods, before calculation of the ‘cell survival ratio’, relative to the number of colonies in the absence of drug. The panels present the mean values from three independent experiments, together with standard deviations.

We previously showed that inhibition of CUL2^LRR1^ in mouse ES cells blocks the ubiquitylation and disassembly of CMG and causes the helicase and associated replisome partner proteins to persist on chromatin after DNA replication ^17^. This can be visualised on condensed patches of constitutive heterochromatin by spinning disk confocal microscopy. GFP-USP37 co-localises on chromatin with CMG partner proteins such as CTF4 (Figure 1B; 92% of cells with GFP-USP37 on condensed heterochromatin also had co-localising CTF4, n = 90) and CLASPIN (Figure S1D; 95% of cells with GFP-USP37 on condensed heterochromatin also had co-localising CLASPIN, n = 112) under such conditions, further suggesting that USP37 associates with the CMG helicase or its partners, independently of CMG ubiquitylation.

### The Pleckstrin-Homology domain of USP37 binds to the CDC45 subunit of CMG

To investigate how USP37 interacts with components or partners of the CMG helicase, we used AlphaFold Multimer ^41^ to screen for potential interactions between mouse USP37 and all the replisome subunits listed in Appendix Tables S1-S4. This revealed a single hit, corresponding to interaction of a previously uncharacterised Pleckstrin-Homology (PH) domain at the amino-terminus of USP37 (Figure 1C, Figure S3) with the CDC45 helicase subunit, with high confidence and in all five models generated by AlphaFold Multimer (Figure 1D, Figure S4).

The predicted binding of the USP37-PH domain to CDC45 was validated experimentally (Figure 1E, interaction of CDC45 with USP37 1-119), using the yeast two-hybrid assay that provides a simple test for AlphaFold predictions of protein-protein interactions ^42^. Subsequently, we purified recombinant mouse USP37 from budding yeast cells (Figure S5) and found that it interacted with recombinant CMG in vitro (Figure 1F, compare lanes 6-7 with lane 8), dependent upon the USP37-PH domain (Figure 1F, lane 10, Figure S5A). Moreover, mutation of conserved sites in the USP37-PH that are predicted to bind to CDC45 (Figure 1D, Figure S3D) was sufficient to prevent the two factors from interacting in the yeast two hybrid assay (Figure 1E, 10A mutations in USP37 1-119). Correspondingly, USP37-10A was unable to interact with the CMG helicase both *in vitro* (Figure S5A, Figure 1F, lane 9) and in mouse ES cells (Figure 1G, compare lanes 6-8). These findings indicated that the USP37-PH domain couples the USP37 deubiquitylase to the CMG helicase via CDC45.

The predicted complex between CDC45 and the USP37-PH domain can be docked onto the cryo-electron-microscopy structure of the human replisome bound to CUL2^LRR1^ ^25^, without clashing with other replisome interactions (Figure 1H). In full length USP37, the PH-domain is connected to the catalytic domain of USP37 by a long disordered linker of around 245 amino acids (Figure S3). This suggests that the USP37 deubiquitylase is recruited to the CMG helicase at DNA replication forks by the USP37-PH domain, but then has considerable spatial flexibility to counteract the action of replisome-associated ubiquitin ligases such as CUL2^LRR1^ and TRAIP.

### Mutations displacing USP37 from the CMG helicase make cells sensitive to DNA replication stress

To investigate the functional significance of the USP37-PH domain in mouse ES cells, we first used CRISPR-Cas9 to impair USP37 expression by making small deletions in the first coding exon of the *Usp37* gene (Figure S2C-I), as a prelude to rescuing such deleted clones with expression of either wild type USP37 or USP37-10A. Large-scale CRISPR screens in human cells indicated that *USP37* is a common essential gene ^43^, and deletion of mouse *Usp37* is embryonic lethal in vivo^44^. However, small frame-shift deletions at the start of the *Usp37* coding sequence are viable, likely due to expression of a very low level of catalytically active protein from internal in-frame initiator codons (Figure S2D). Correspondingly, frame-shift mutations within the USP37 catalytic domain, produced by targeting CRISPR-Cas9 to Exon 11 or Exon 12 of *Usp37*, are inviable and all viable clones were found to express approximately full-length USP37 protein with very small in-frame deletions (Figures S6-S7). These data indicate that *Usp37* is an essential gene in mouse ES cells, suggesting that viable frame-shift deletions at the start of the coding sequence (hereafter referred to as ‘*Usp37^hypo^*’) represent hypomorphic alleles that provide useful tools for functional analysis.

Depletion of USP37 sensitises human cells not only to agents that inhibit DNA synthesis at replication forks ^34,38,45^, but also to the anti-cancer drug camptothecin ^39^, which traps Topoisomerase on DNA and leads to topological stress in front of replication forks. Correspondingly, *Usp37^hypo^* mouse ES cells are highly sensitive to hydroxyurea that depletes deoxynucleotides, aphidicolin that inhibits DNA polymerases and also to camptothecin (Figure 1I, compare Control and *Usp37^hypo^*). The sensitivity of *Usp37^hypo^* to all three forms of DNA replication stress can be rescued by expression of wild type USP37 from the *Rosa26* ‘safe-haven’ locus (Figure 1I, *Usp37^hypo^* + GFP-USP37), but not by equivalent expression of USP37-10A (Figure 1I, *Usp37^hypo^* + GFP-USP37-10A). These findings indicate that the PH domain of USP37 is important to protect mammalian cells from a range of DNA replication stresses, including DNA synthesis defects and topological stress.

### The sensitivity of *Usp37^hypo^* cells to topological stress is suppressed by removal of the TRAIP ubiquitin ligase

The importance of the USP37-PH domain during DNA replication stress suggested that USP37 might act at replication forks in mammalian cells, to counteract ubiquitin ligases that associate with the replisome, such as CUL2^LRR1^ and TRAIP. In that case, inactivation of either CUL2^LRR1^ or TRAIP might suppress the sensitivity of *Usp37^hypo^* cells to agents that either cause DNA synthesis defects or that induce topological stress during chromosome replication.

To begin to test this idea, we used CRISPR-Cas9 to generate *Usp37^hypo^ TraipΔ* mouse ES cells (Figure 2). The *Usp37^hypo^ TraipΔ* double mutant is just as sensitive as *Usp37^hypo^* to DNA synthesis defects produced by treating cells with hydroxyurea (Figure 2A-B) or aphidicolin (Figure 2C-D). Strikingly, however, deletion of *Traip* suppressed the sensitivity of *Usp37^hypo^* cells to camptothecin (Figure 2E-F). These data indicate that the USP37 deubiquitylase counteracts the action of the TRAIP ubiquitin ligase at DNA replication forks, when cells experience topological stress during chromosome replication. However, the data also suggest that USP37 might be needed to counteract a different ligase (or ligases) when cells experience DNA synthesis defects in response to agents such as hydroxyurea or aphidicolin.

**Figure 2.**
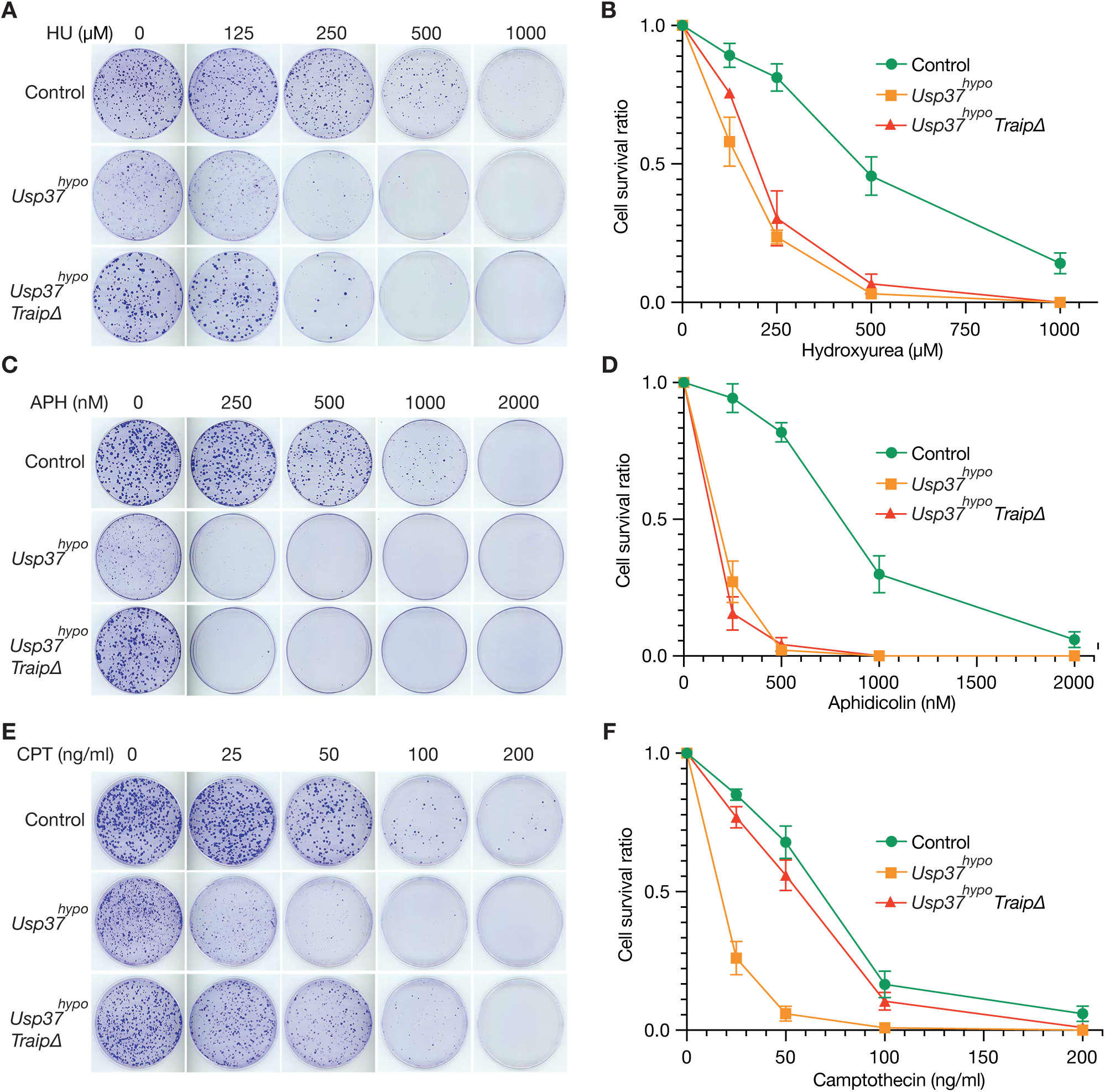
The sensitivity of *Usp37^hypo^* cells to topological stress is suppressed by removal of the TRAIP ubiquitin ligase. (**A**) Mouse ES cells of the indicated genotypes were treated with the indicated concentrations of hydroxyurea (HU), as described above for Figure 1I. (**B**) Quantification of the data as in Figure 1I. The panel presents the mean values from three independent experiments, together with standard deviations. (**C**) Analogous experiment to (A) but with the indicated concentrations of aphidicolin (APH). (**D**) Quantification of the data as in (B). (**E**) Analogous experiment to (A) but with the indicated concentrations of camptothecin (CPT). (**F**) Quantification of the data as in (B).

### USP37 counteracts CMG ubiquitylation by the CUL2^LRR1^ ubiquitin ligase

We previously showed that CUL2^LRR1^ ubiquitylates the CMG helicase on its MCM7 subunit in mammalian cells ^16,17^, leading rapidly to disassembly by the p97 unfoldase, consistent with findings in *Xenopus laevis* egg extracts ^14^ and the early embryo of *Caenorhabditis elegans* ^15^. When control and *Usp37^hypo^* cells were treated with a small-molecule inhibitor of p97, the ubiquitin chains on CMG-MCM7 became longer in the absence of USP37 (Figure 3A, compare lanes 7-8). This effect could be rescued by expression of wild type USP37 at the *Rosa26* locus in *Usp37^hypo^* cells (Figure 3B, compare lanes 6-7), but not by expression of USP37 with mutation of a conserved catalytic residue ^37^ (Figure 3B, USP37-C350A, lane 8), which is unable to deubiquitylate CMG in vitro (Figure 3C), or rescue the sensitivity of *Usp37^hypo^* to DNA replication stress (Figure 3D). These findings indicate that USP37 deubiquitylase activity counteracts CMG helicase ubiquitylation and protects cell viability during DNA replication stress.

**Figure 3.**
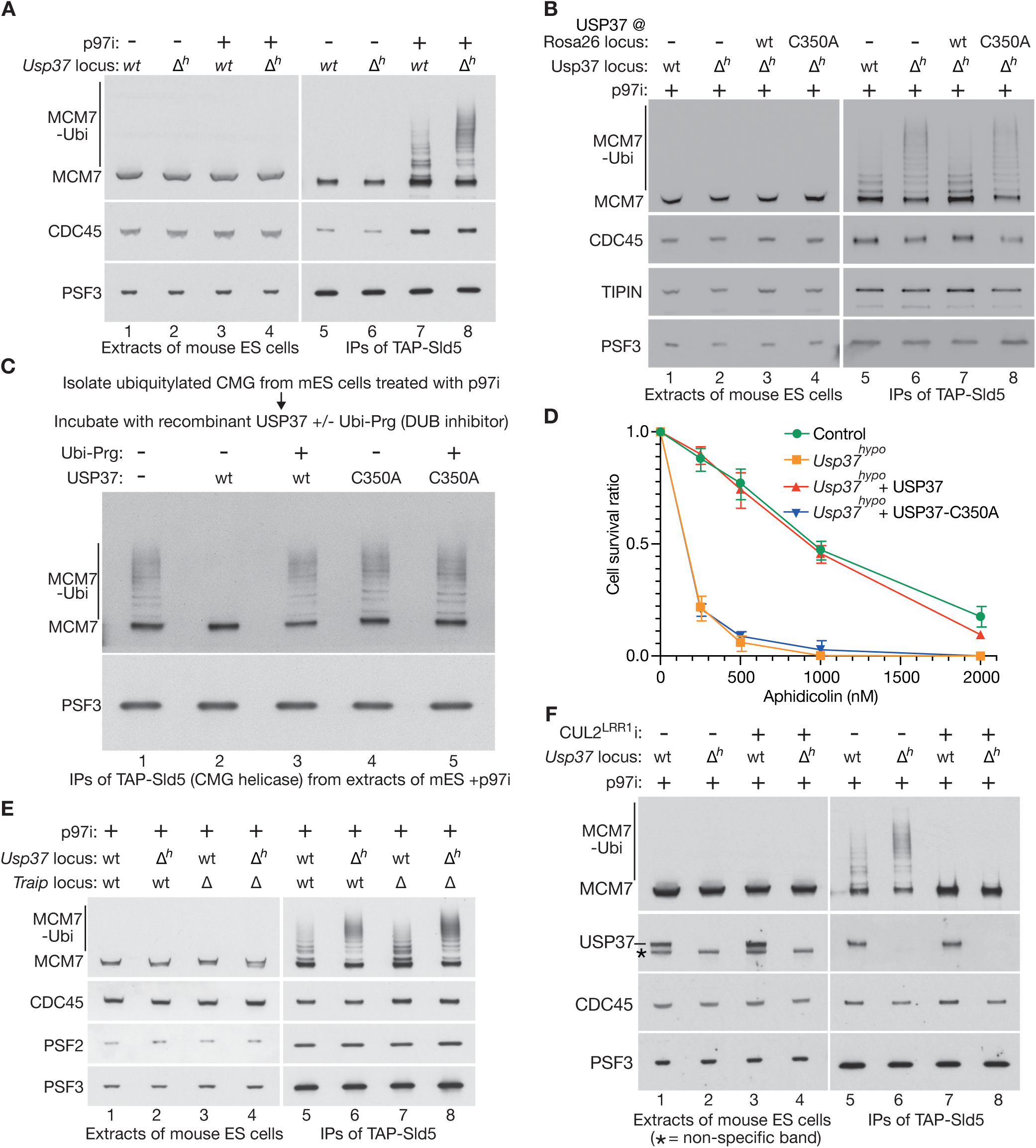
USP37 deubiquitylase activity counteracts CMG ubiquitylation by CUL2^LRR1^. (**A-B**) Mouse ES cells of the indicated genotype were treated with 5 µM CB-5083 (p97 inhibitor = p97i) as shown. Extracts were incubated with IgG-coated magnetic beads and the immunoprecipitated proteins monitored by immunoblotting. ‘*Δh*’ = *Usp37^hypo^*. (**C**) Ubiquitylated CMG helicase was isolated from *TAP-Sld5* mouse ES cells treated with 5 µM CB-5083 (p97i). The immunoprecipitates were then treated for 1 hour at 24 °C with recombinant USP37 or USP37-C350A as described in STAR methods, in the presence or absence of the deubiquitylase (DUB) inhibitor Ubi-Prg (Propargylated Ubiquitin) as indicated. The reactions were then monitored by SDS-PAGE and immunoblotting. (**D**) Equivalent experiment to (Figure 1I) but also including *Usp37^hypo^* cells rescued by either wild type USP37 (*Usp37^hypo^ +USP37*) or the catalytically dead USP37-C350A allele (*Usp37^hypo^ +USP37-C350A*). (**E**) Mouse ES cells of the indicated genotype were processed as in (A). (**F**) CUL2^LRR1^ was inhibited as shown in the indicated mouse ES cells, as described in STAR methods, before inhibition of p97. Samples were then processed as above.

To determine which ubiquitin ligase is responsible for enhanced CMG ubiquitylation in *Usp37^hypo^* cells, we first compared the effect of inhibiting p97 in control, *Usp37^hypo^*, *TraipΔ* or *Usp37^hypo^ TraipΔ* cells. Deletion of Traip did not impair CMG ubiquitylation in response to p97 inhibition (Figure 3E, compare lanes 5 and 7), under conditions where the CMG helicase persisted on chromatin after DNA replication termination, and entry into mitosis was impaired. Moreover, the ubiquitylation of the CMG-MCM7 subunit was enhanced to a similar degree in *Usp37^hypo^* (Figure 3E, lane 6) and *Usp37^hypo^ TraipΔ* cells (Figure 3E, lane 8) after treatment with p97 inhibitor.

In contrast, CMG ubiquitylation was completely abrogated under such conditions by inhibition of CUL2^LRR1^, both in the control and in *Usp37^hypo^* cells (Figure 3F, lanes 7-8). These findings indicated that USP37 counteracts CMG ubiquitylation by CUL2^LRR1^ in mammalian cells.

### Depletion of LRR1 suppresses the sensitivity of *Usp37^hypo^* cells to DNA synthesis defects

To test whether CUL2^LRR1^ activity was responsible for the sensitivity of *Usp37^hypo^* cells to defects in DNA synthesis, we investigated the functional consequences of inactivating CUL2^LRR1^ in *Usp37^hypo^* cells. LRR1 is essential for viability in human cells ^46,47^ and during mouse development ^44^. Therefore, we generated mouse ES cells in which the BromoTag degron ^48^ was introduced before the initiator codon of the *Lrr1* coding sequence (Figure S8A). This enabled the rapid degradation of BromoTag-LRR1 when cells were treated with AGB1 (Figure S8B), which is a highly selective and potent Proteolysis Targeting Chimera (PROTAC) that links the BromoTag degron cassette to the CUL2^VHL^ ubiquitin ligase ^48^. Degradation of BromoTag-LRR1 is lethal in mouse ES cells (Figure S8C), consistent with the essential nature of LRR1 in mice and human cells ^44,46,47^.

To test whether depletion of LRR1 can suppress *Usp37^hypo^* phenotypes, we initially tried to degrade BromoTag-LRR1, then expose cells to DNA replication stress, and finally remove AGB1 to allow re-expression of the fusion protein. However, efficient depletion of BromoTag-LRR1 is lethal, even in the absence of DNA replication stress. Subsequently, therefore, we took advantage of the fact that the level of BromoTag-LRR1 degradation can be controlled by titration of AGB1 (Figure S8D-E). This allowed us to screen for a ‘sweet spot’ in which the residual level of BromoTag-LRR1 is sufficient to support cell viability, but is too low to cause problems at stalled DNA replication forks in the absence of USP37. In this way, we chose to expose cells to 5 nM AGB1 in all subsequent experiments, leading to an approximately 10-fold depletion of BromoTag-LRR1 (Figure S8D-E, lane 4), without causing lethality (Figure S8F-G).

When grown in the presence of camptothecin and 5 nM AGB1, the behaviour of control and *BromoTag-Lrr1* cells was very similar to control cells treated with camptothecin but lacking PROTAC (compare Figure 4A with Figure 1I right panel). As expected, *Usp37^hypo^* cells were highly sensitive to camptothecin under such conditions (Figure 4A, *Usp37^hypo^* + 5 nM AGB1) and *Usp37^hypo^ BromoTag-Lrr1* behaved very similarly (Figure 4A, *Usp37^hypo^ BromoTag-Lrr1* + 5 nM AGB1). Therefore, reducing the level of LRR1 did not suppress the sensitivity of *Usp37^hypo^* cells to camptothecin, unlike deletion of *Traip* in *Usp37^hypo^* cells (Figure 2E-F).

**Figure 4.**
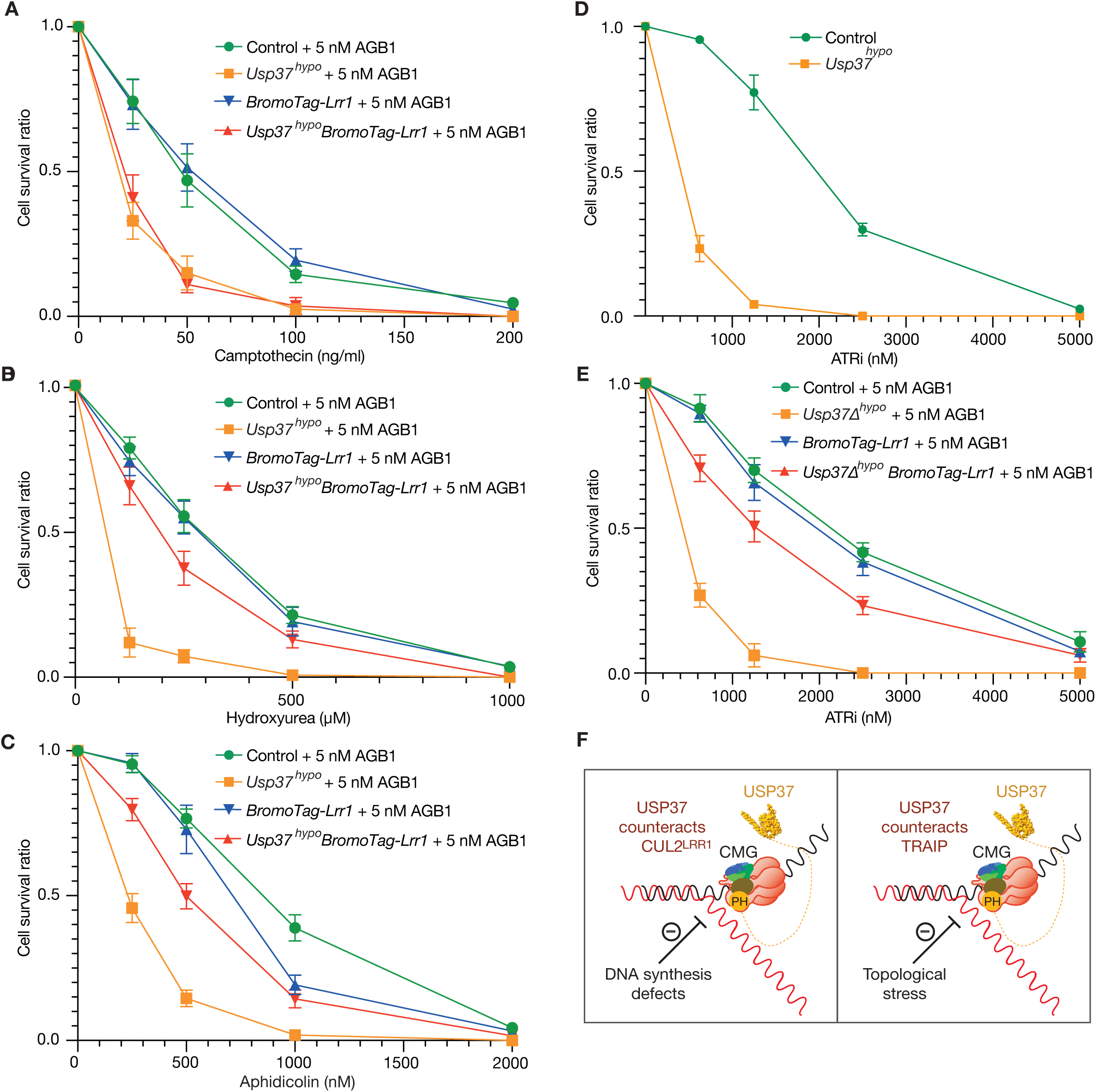
Depletion of LRR1 suppresses the sensitivity of *Usp37^hypo^* cells to DNA synthesis defects and ATR kinase inhibitors. (**A**) Mouse ES cells of the indicated genotypes were treated with increasing concentrations of camptothecin as shown, in the presence of 5 nM AGB1 to induce partial depletion of BromoTag-LRR1. Samples were processed and quantified as described above for Figure 1I. The data represent the mean values from three independent experiments, together with standard deviations. (**B**) Analogous experiment to (A) but with the indicated concentrations of Hydroxyurea. (**C**) Analogous experiment to (A) but with the indicated concentrations of Aphidicolin. (**D**) Control mouse ES and *Usp37^hypo^* cells were grown in the presence of the indicated concentrations of the ATR inhibitor AZD6738 (ATRi) for 24 hours, before growth for a further seven days in medium lacking ATRi. Colonies were stained with crystal violet and then quantified as above. (**E**) Analogous experiment to (D) but with mouse ES cells of the indicated genotypes and partial depletion of BromoTag-LRR1 by addition of 5 nM AGB1 to the culture medium. Data were quantified as above. (**F**) Model illustrating the action of the USP37 deubiquitylase at DNA replication forks, in opposition to the CUL2^LRR1^ ubiquitin ligase (in response to DNA synthesis defects) and the TRAIP ubiquitin ligase (during topological stress).

In contrast, depletion of BromoTag-LRR1 dramatically suppressed the sensitivity of *Usp37^hypo^* cells to hydroxyurea (Figure 4B, compare *Usp37^hypo^* + 5 nM AGB1 with *Usp37^hypo^ BromoTag-Lrr1* + 5 nM AGB1). Moreover, depletion of BromoTag-LRR1 also strongly suppressed the sensitivity of *Usp37^hypo^* cells to aphidicolin (Figure 4C, compare *Usp37^hypo^* + 5 nM AGB1 with *Usp37^hypo^ BromoTag-Lrr1* + 5 nM AGB1). These data indicate that USP37 protects cells during DNA synthesis defects by counteracting the untimely action of CUL2^LRR1^. Furthermore, the above findings indicate that USP37 protects cells from different replisome-coupled ubiquitin ligases in response to topological stress (TRAIP) and DNA synthesis defects (CUL2^LRR1^).

### Dependency of *Usp37^hypo^* cells on the ATR checkpoint kinase is suppressed by LRR1 depletion

In human cells, USP37 depletion by siRNA or CRISPR-Cas9 leads to increased stalling of DNA replication forks and some degree of genome instability, even in the absence of agents that cause exogenous DNA replication stress or topological stress ^33,38,45^. For this reason, depletion of USP37 causes sensitivity to ATR checkpoint kinase inhibitors ^39^, which kill cells with endogenous sources of DNA replication stress and are currently in clinical trials as an anti-cancer treatment ^49^.

Consistent with the human data, *Usp37^hypo^* sensitises mouse ES cells to ATR inhibition (Figure 4D), regardless of the presence or absence of the AGB1 PROTAC (compare Figure 4D-E). In the presence of 5 nM AGB1, *BromoTag-Lrr1* cells behaved very similarly to control cells treated with ATR inhibitor (Figure 4E). However, depletion of BromoTag-LRR1 strongly suppressed the sensitivity of *Usp37^hypo^* cells to ATR inhibition (Figure 4E, *Usp37^hypo^ BromoTag-Lrr1*). These data indicate that USP37 is important to counteract the action of CUL2^LRR1^ in mammalian cells, even in the absence of exogenous sources of DNA replication stress.

## Discussion

The mammalian USP37 deubiquitylase has been reported to have a diverse range of targets ^33,36,37,50–55^, yet the major phenotypes of cells lacking USP37 are sensitivity to agents that cause DNA replication stress ^34,39^, increased stalling of DNA replication forks ^33,34,45^, defects in the DNA damage response ^35,36,38^ and defects in resolving sister chromatids during mitosis ^37^. The data presented in this manuscript indicate that the PH domain at the amino terminus of USP37 is essential to protect cells from DNA replication defects and topological stress (Figure 1I), serving to tether the USP37 deubiquitylase to DNA replication forks by binding to the CDC45 subunit of the CMG helicase (Figure 1D-H). It is likely that USP37 can access a range of potential substrates within the vicinity of CMG, as well as counteracting ubiquitylation of the CMG helicase itself, due to the unusually long disordered linker that connects the PH domain of USP37 to the catalytic domain (Figure 1H, Figure S3).

Our data support a model whereby USP37 deubiquitylase activity has two important and related roles at DNA replication forks (Figure 4F). Firstly, USP37 protects cells in response to DNA synthesis defects by counteracting the function of the CUL2^LRR1^ ubiquitin ligase (Figure 3, Figure 4B-C). CMG ubiquitylation by CUL2^LRR1^ is enhanced in the absence of USP37 (Figure 3F), and USP37 can reverse CUL2^LRR1^-mediated CMG ubiquitylation in vitro (Figure 3C), indicating that the deubiquitylase is a direct regulator of the CMG helicase that opposes the action of CUL2^LRR1^ (Figure 4F, left panel). However, CUL2^LRR1^ rapidly induces CMG ubiquitylation and disassembly during DNA replication termination, despite the presence of USP37. Moreover, the ability of CUL2^LRR1^ to ubiquitylate CMG is sterically blocked at DNA replication forks, independently of USP37, by the lagging-strand DNA template that blocks a key contact site between LRR1 and the helicase^25^. We propose that USP37 is important to counteract CUL2^LRR1^ function at stalled DNA forks in mammalian cells, in response to defects in DNA synthesis. Subsequently, the bypass of two replisomes during DNA replication termination removes the steric block to CUL2^LRR1^ and unleashes the ligase to promote efficient CMG ubiquitylation, thereby outcompeting USP37.

The second important role for USP37 is to counteract the TRAIP ubiquitin ligase in response to topological stress (Figure 2E-F, Figure 4F right panel). We note that replication forks are particularly vulnerable to topological stress when they converge during termination ^8,56^. This might increase the risk that TRAIP acts in trans at converging replisomes to mediate replisome ubiquitylation and disassembly, before helicase bypass and the completion of DNA synthesis has occurred, thereby leaving behind small unreplicated segments of the genome. Tethering of USP37 to the CMG helicase would mitigate this risk by counteracting TRAIP, thereby explaining the sensitivity of *Usp37^hypo^* cells to camptothecin. Whether topological stress leads to such events in untreated cells remains to be explored in future studies, but camptothecin derivatives are used in the treatment of a wide variety of solid tumours in humans ^57,58^, raising the possibility that USP37 inhibitors might have a synergistic role of potential therapeutic benefit.

Whereas CUL2^LRR1^ and TRAIP are broadly conserved in diverse animal species, USP37 is present in vertebrates but appears to be missing in many other groups of metazoans. It will be interesting in future studies to explore whether stalled DNA replication forks are particularly vulnerable to the untimely action of CUL2^LRR1^ and TRAIP in vertebrate species, either in response to DNA replication defects or as a result of topological stress. For example, the processing of stalled DNA replication forks by factors such as RAD51 ^59^ or fork-reversal enzymes ^60^ might transiently break the association of the helicase with the excluded parental DNA strand, thereby briefly exposing CMG to CUL2^LRR1^ at replication forks. Under such conditions, USP37 might have evolved or been retained in order to counteract CUL2^LRR1^ and preserve genome integrity.

Cells lacking USP37 function have a heightened dependency on the ATR checkpoint kinase that preserves the integrity of stalled DNA replication forks (Figure 4D, ^39^). Inhibitors of ATR are currently in clinical trials as anti-cancer agents ^49^, and inhibition of the USP37 deubiquitylase would be predicted to synergise with drugs inhibiting ATR. Our data demonstrate that depletion of LRR1 represents a potential resistance mechanism that could suppress the sensitivity of cells to the combined inhibition of USP37 and ATR (Figure 4E). These data further suggest that counteracting CUL2^LRR1^ was a major driver for the evolution of USP37 in vertebrate cells.

Over-expression of USP37 has been reported in a number of human cancers and correlates with with poor prognosis ^54,61^. The importance of USP37 deubiquitylase activity for cell survival when DNA replication is defective (Figure 3D), and the synthetic lethal effects of inactivating USP37 in combination with anti-cancer agents such as ATR inhibitors or camptothecin, make it important to explore the impact of USP37 inhibition in tumours with high levels of endogenous DNA replication stress.

## Acknowledgements

We thank Axel Knebel for propargylated ubiquitin, MRC PPU Reagents and Services (https://mrcppureagents.dundee.ac.uk) for antibody production and Johannes Walter, Steve Jackson, Nicholas Brown, Jean Cook and Michael Emanuele for discussing unpublished data. We are grateful for the support of the Medical Research Council (core grants MC_UU_12016/13 to KL) and Cancer Research UK (Programme Grant C578/A24558 and PhD studentship C578/A25671 to KL). Materials generated in this study are listed in STAR Methods and are available from MRC PPU Reagents and Services (https://mrcppureagents.dundee.ac.uk) or upon request.

## Author Contributions

All experiments were performed by FV, except for Figure S1B-C and the data summarised in Supplementary Tables 3-4, which were performed by JA. KL supervised the project and wrote the manuscript in collaboration with FV and JA.

## MATERIALS AND METHODS

### Materials availability

Materials generated in this study are available from MRC PPU Reagents and Services (https://mrcppureagents.dundee.ac.uk) or upon request.

### Data and code availability

- Mass spectrometry data have been deposited at the ProteomeExchange Consortium via the PRIDE partner repository database ^63^ and will be made publicly available by the date of publication. Accession numbers will be listed in Table S5. Microscopy data reported in this paper will be shared by the lead contact upon request.
- This paper does not report original code.
- Any additional information required to reanalyze the data reported in this paper is available from the lead contact upon request.

### Growth of Mouse Embryonic Stem cells

Mouse ES cells (E14tg2A) were maintained in Dulbecco’s Modified Eagle Medium (DMEM, ThermoFisher Scientific, 11960044) supplemented with 10% Foetal Bovine Serum (FBS, FCS-SA/500, LabTech), 5% KnockOut Serum Replacement (ThermoFisher Scientific, 10828028), 2 mM L-Glutamine (ThermoFisher Scientific, 25030081), 100 U/ml Penicillin–Streptomycin (ThermoFisher Scientific, 15140122), 1 mM Sodium Pyruvate (ThermoFisher Scientific, 11360070), a mixture of seven non-essential amino acids (ThermoFisher Scientific, 11140050), 0.05 mM β-mercaptoethanol (Sigma-Aldrich, M6250) and 0.1 μg/ml Leukaemia Inhibitory Factor (MRC PPU Reagents and Services, DU1715). Cells were grown at 37°C in a humidified atmosphere of 5% CO_2_, 95% air. For passaging, cells were released from dishes using 0.05% Trypsin–EDTA (ThermoFisher Scientific, 25300054). Degradation of BromoTag-LRR1 was induced by addition of indicated concentrations of the PROTAC AGB1 (Tocris, 7686).

To inhibit the p97 ATPase, cells were treated with 5 µM CB-5083 (S8101, Selleckchem) for 3 hours before harvesting. To inhibit the activity of cullin ligases, 5 µM MLN4924 (A-1139, Activebiochem) was added to the culture medium for 5 hours before harvesting. For the experiments in Figure 1B, Figure 3F and Figure S1D, CUL2^LRR1^ was inhibited by transfection with *Lrr1* siRNA as described below, combined with 5 µM MLN4924 for the last five hours.

### Generation of plasmids expressing gRNAs for CRISPR-Cas9 genome editing

To facilitate genome editing by the *Streptococcus pyogenes* Cas9-D10A nickase, a pair of guide RNAs (gRNAs) was designed using “CRISPR Finder” (https://wge.stemcell.sanger.ac.uk//find_crisprs), to target a specific site in the mouse genome. Each gRNA contains 20 nucleotides of homology, which in the genome is located immediately upstream of a three nucleotide “Protospacer Adjacent Motif” (PAM) sequence that is required for cleavage by Cas9-D10A. Two annealed oligonucleotides containing the homology region were phosphorylated by T4 polynucleotide kinase (New England Biolabs, M201) and cloned into the vectors pX335 and pKN7 ^17^ after digestion by the Type IIS restriction enzyme BbsI (New England Biolabs, R3539) as previously described ^64^.

### Generation of donor vectors to introduce tags into mammalian genes by CRISPR-Cas9

Donor vectors for the introduction of purification tags, fluorescent proteins or degron cassettes into genomic loci in mammalian genomes, were made using the previously described “mammalian toolkit” using Golden Gate cloning ^65^ and contained ∼800-1,000 bp homology to the target locus and a hygromycin selection marker. Silent mutations were introduced to ensure that neither of the two gRNAs would be able to anneal with the target locus after successful genome editing. Donor vectors to be introduced at the *Rosa26* locus contained the EF1-a promoter to express the chosen coding sequence.

For molecular biology purposes PCRs were performed with PrimeStar Hot Start DNA polymerase (Takara, R010A). All new constructs were verified by Sanger sequencing. Site specific mutagenesis was performed using PrimeStar Hot Start DNA polymerase, according to the manufacturer’s protocol.

### Buffers

#### Yeast Lysis buffer

25 mM HEPES KOH pH 7.6, 0.15 M NaCl, 10% glycerol, 5 mM MgOAc, 1 mM DTT (dithiothreitol).

#### Yeast Wash buffer

50 mM HEPES KOH pH 7.6, 0.15 M NaCl, 10% glycerol, 1 mM DTT, 0.02% (v/v) IGEPAL CA-630.

#### FLAG elution buffer

50 mM HEPES KOH pH 7.6, 0.15 M NaCl, 10% glycerol, 1 mM DTT.

#### Gel filtration buffer

25 mM HEPES KOH pH 7.6, 0.15 M NaCl and 1 mM DTT.

#### Protein binding buffer

25 mM Hepes-KOH pH 7.6, 200 mM KOAC, 0.02% IGEPAL CA-630, 10 mM MgOAc, 1 mM DTT and 0.1 mg/ml BSA.

#### DUB buffer

100mM HEPES-KOH pH 7.9, 100mM Potassium Acetate, 10 mM Magnesium Acetate, 0.02 % IGEPAL CA-630, 1mM DTT.

#### mES lysis buffer

100 mM Hepes-KOH pH 7.9, 100 mM potassium acetate, 10 mM magnesium acetate, 2 mM EDTA, 10% glycerol, 0.1% Triton X-100, 2 mM sodium fluoride, 2 mM sodium β-glycerophosphate pentahydrate, 1 mM dithiothreitol.

#### mES wash buffer

100 mM Hepes-KOH pH 7.9, 100 mM potassium acetate, 10 mM magnesium acetate, 2 mM EDTA, 10% glycerol, 0.1% IGEPAL CA-630, 2 mM sodium fluoride, 2 mM sodium β-glycerophosphate pentahydrate, 1 mM dithiothreitol.

### Protease inhibitors

In this study, ‘Protease inhibitors’ corresponds to a combination of 1X ‘Roche cOmplete EDTA-free protease inhibitor Cocktail’ and 1% ‘Protease Inhibitor Cocktail’ (Sigma-Aldrich, P8215). One tablet of ‘Roche cOmplete EDTA-free protease inhibitor Cocktail’ (Roche, 11873580001) was dissolved in 1 ml of water to make a 20X stock.

### USP37 expression and purification from budding yeast

The *Saccharomyces cerevisiae* strains used in this study are shown in Table S5. Yeast cells were grown at 30°C in YP medium (1 % Yeast Extract, 21275, Becton Dickinson; 2 % bacteriological peptone, LP0037B, Oxoid) supplemented with 2 % Raffinose. To induce protein expression, a 3-litre exponential culture was grown to 2-3 x 10^7^ cells/ml before addition of Galactose to a final concentration of 2 % and further incubation for 4 h. Cells were collected by centrifugation and washed once with ‘Yeast lysis buffer’ without protease inhibitors. The cell pellet (∼ 10 g) was then resuspended in 0.3 volumes of lysis buffer containing protease inhibitors. The resulting suspension was frozen dropwise in liquid nitrogen and stored at - 70°C. Subsequently, the frozen yeast cells were ground in the presence of liquid nitrogen, using a SPEX CertiPrep 6850 Freezer/Mill with 3 cycles of 2 minutes at a rate of 15. The resulting powder was then stored at -70 °C, before thawing in Yeast lysis buffer with protease inhibitors. Insoluble material was removed by centrifugation (235,000 g, 4°C, 1 h) and the supernatant was mixed with 1 mL anti-FLAG M2 affinity gel (Sigma-Aldrich) at 4°C for 90 min.

The resin was collected and washed with ‘Yeast Wash Buffer’. USP37 was eluted in 1 column volume of ‘FLAG elution buffer’ with 0.5 mg / ml 3FLAG peptide, then 1 column volume of FLAG elution buffer with 0.25 mg / ml 3FLAG peptide.

The USP37 containing fractions were pooled, concentrated and separated on a 24 mL Superdex 200 column in ‘Gel filtration buffer’. Subsequently, USP37 containing fractions were again pooled and concentrated, before being aliquoted and snap frozen.

### Preparation of antibody-coated magnetic beads

A 300 mg aliquot of Dynabeads M-270 Epoxy (ThermoFisher Scientific, 14302D) was resuspended in 10 ml of dimethylformamide, producing an activated slurry of magnetic beads. Subsequently, 425 µl (∼ 1.4 x 10^9^ beads) was washed twice with 1 ml of 1 M NaPO_3_ (pH 7.4). Beads were incubated with 300 µl of 3 M (NH4)_2_SO_4_, 300 µg of anti-FLAG M2 antibody, and sufficient 1 M NaPO_3_ (pH 7.4) to make a total volume of 900 µl. After incubation at 4 °C for two days with rotation, the beads were treated as follows: four washes with 1 ml PBS, 10 minutes in 1 ml PBS containing 0.5 % (v/v) IGEPAL CA-630 at room temperature with rotation, five minutes in 1 ml PBS containing 5 mg / ml BSA at room temperature with rotation, followed by one wash with 1 ml PBS containing 5 mg / ml BSA. Finally, the beads were resuspended in 900 µl PBS containing 5 mg / ml BSA and stored at 4°C.

### In vitro binding assay for mouse CMG and USP37

For the experiment in Fig 1F, 100 nM purified recombinant mouse USP37 and 20 nM CMG ^65^ were incubated in ‘Protein Binding Buffer’ for 30 minutes on ice. Subsequently, 5 μl of magnetic beads coupled to anti-FLAG M2 antibody was added to the protein mixture and incubated for 1 h on ice with occasional mixing. After two washes with the ‘Protein Binding Buffer’, the bound proteins were eluted from the beads with 15 μl of 1× Laemmli buffer by heating at 95°C for 5 minutes. Proteins were resolved in a 4–12% Bis-Tris NuPAGE gel (Life Technologies, NP0301) in 1× MOPS buffer and then detected by immunoblotting.

### In vitro deubiquitylation assay

For the experiment in Figure 3C, ubiquitylated CMG was isolated as described above by immunoprecipitation of TAP-SLD5 from mouse ES cells treated with 5 µM CB-5083. The purified complexes were then treated for 1 hour at 24 °C with 0.1 µM USP37 or USP37-C350A in ‘DUB buffer’, with agitation at 1400 rpm on an Eppendorf Thermomixer F1.5. The reactions were stopped by addition of 20 μl of 3X Laemmli buffer and then heated for 5 minutes at 95 °C, before analysis by immunoblotting.

### SDS-PAGE and Immunoblotting

Proteins were separated in 4–12% Bis-Tris NuPAGE gels (Life Technologies, NP0301) using 1× MOPS buffer. Alternatively, ubiquitylated forms of MCM7 were resolved in 3-8% Tris-Acetate gels (Life Technologies, WG1602BOX) with 1X Tris-Acetate SDS Running Buffer. Subsequently, samples were either stained with InstantBlue (Expedeon, ISB1L), or transferred to a nitrocellulose membrane and detected by immunoblotting using the iBlot Dry Transfer System (ThermoFisher). Primary antibodies used for protein detection in this study are described in Table S5. Secondary antibodies are also listed in Table S5 and corresponded to horseradish peroxidase conjugates of anti-sheep IgG from donkey (Sigma-Aldrich, A3415), or anti-mouse IgG from goat (Sigma-Aldrich, A4416). Chemiluminescent signals were detected on Hyperfilm ECL (Cytiva, 28906837, 28906839) or the AZURE 300 imaging system (Azure Biosystems), using the ECL Western Blotting Detection Reagent (Cytiva, RPN2124).

### Transfection of mouse ES cells and selection of clones

Cells were transfected with plasmids that expressed the Cas9-D10A “nickase”, gRNAs specific for the target locus (one for *Rosa26*, two in all other cases), and the Puromycin resistance gene. Donor vectors were included as required.

Aliquots of 1.0 × 10^6^ cells were centrifuged at 350 *g* for 5 minutes before resuspension in a mixture of 200 μl DMEM, 15 μl of 1 mg/ml linear Polyethylenimine (PEI; Polysciences, Inc, 24765-2) and 1–2 μg of each plasmid. The cells were then mixed gently by pipetting and incubated at room temperature for 30 minutes, before transfer to a single well of a 6-well plate precoated with 0.1% gelatin (Sigma, G1890) and containing 2 ml of complete DMEM medium. After incubation for 24 h, cells were subjected to two 24-h rounds of selection with fresh medium containing 2 μg/ml Puromycin (ThermoFisher Scientific, A1113802), followed by 1–2 weeks of selection with fresh medium containing 200–500 μg/ml Hygromycin (Invitrogen, ant-hg-5). The surviving cells were released from the well with 0.05% Trypsin–EDTA (ThermoFisher Scientific, 25300054), diluted from 10 to 10,000 times with complete DMEM containing Hygromycin, and plated on 10 cm plates precoated with 0.1% gelatin. Subsequently, single colonies were picked and transferred to one well of a 6-well plate precoated with 0.1% gelatin. After 6–10 days, viable clones were expanded and monitored by immunoblotting, PCR and DNA sequencing of the target locus.

### Transfection of siRNA into mouse ES cells

Lipofectamine RNAiMAX (13778, ThermoFisher) was used to introduce siRNA into mouse ES cells, according to the manufacturer’s protocol. Details of the *Lrr1* and control siRNAs are given in Table S5 and were all transfected at a final concentration of 25 nM. After 24 hours, the transfection was repeated, and incubation continued for a further 24 hours.

### PCR and DNA sequence analysis for genotyping of mouse ES cells

Cells from a 10 cm plate were released as described above. One third of the cells were pelleted and resuspended in 100 μl of 50 mM NaOH, before heating at 95°C for 15 minutes. The pH was then neutralised by addition of 11 μl of 1 M Tris– HCl (pH 6.9) and 0.5 μl used as a source of genomic DNA in 20 μl PCRs with PrimeStar Hot Start DNA polymerase (Takara Bio, R010A). The amplified DNA was subcloned with StrataClone Blunt PCR cloning kit (Agilent Technologies, 240207) and sequenced with M13 Forward, M13 Reverse primers, together with primers specific to the target locus as necessary.

### Cell survival assays for mouse ES cells

For each sample, 500 cells (800 cells for *Usp37hypo* lines) were plated into a 10 cm petri dish with the indicated concentration of drug. After 24 hours, the medium was changed with fresh medium lacking the drug, and incubation proceeded for a further 7 days. The cells were then washed with PBS and fixed with methanol. The fixed cells were stained with 0.5 % crystal violet solution (HT90132, Sigma Aldrich), and images of the plates were then captured with a scanner. Cell colonies were counted with the aid of the ImageJ software (National Institutes of Health) as follows. The image was first converted to 8-bit, then a circle was drawn around the edge of the dish with the oval ‘Region of Interest’ tool. Under the “Edit” tab, the command “Clear Outside” was used to remove the area outside of the oval from the analysis area. The threshold, located in the “Image” tab under “Adjust”, was set to make sure colonies were highlighted as red spots with a black background. Under “Process” and “Binary”, the tool “Watershed” was then used to split merging colonies. Finally, colonies were counted using the “Analyse Particles” command. Data were then analysed with GraphPad Prism software (GraphPad Software).

### Preparation of denatured extracts of mouse ES cells with trichloroacetic acid (TCA)

To monitor BromoTag-LRR1 in the experiments in Figures S8D-E, 1 × 10^6^ cells were seeded per well in 6-well plates. After treatment with PROTAC as indicated, cells were released from the wells with 0.05% Trypsin–EDTA, washed with PBS and resuspended in 100 μl of 20% TCA. The suspension was then mixed by vortexing for 1 min and centrifugated for 10 minutes at 845 *g* at room temperature. The supernatant was then discarded, and the pellet resuspended in 75 μl of Laemmli buffer adjusted to 150 mM Tris-base, followed by addition of 25 μl of 4× NuPAGE LDS sample buffer (ThermoFisher Scientific, NP0007) before incubation at 95°C for 10 minutes.

### Preparation of native extracts of mouse ES cells and immunoprecipitation

Cells were seeded on 15 cm plates at 1.5–2 × 10^7^ cells per dish. After 48 h at 37°C, cells were released from the plates by incubation with PBS containing 1 mM EDTA and 1 mM EGTA for 10 min and harvested by centrifugation. Cells were then resuspended in one volume of ‘mES lysis buffer’ containing protease inhibitors and also 5 μM Propargylated-Ubiquitin (MRC PPU Reagents & Services, DU49003) to inhibit deubiquitylase activity. Chromosomal DNA was digested for 30 minutes at 4°C with 1,600 U/ml of Pierce Universal Nuclease (ThermoFisher Scientific, 88702) before centrifugation at 20,000 *g* for 30 minutes at 4°C.

For immunoprecipitation experiments, cells from two 15 cm dishes cells were utilised. Extracts prepared as above extract were incubated for 90 minutes with either a 15 μl slurry of ChromoTek GFP-Trap magnet Particles M-270 (Proteintech, gtd-100), or a 15 μl slurry of M-270 epoxy beads coupled to rabbit IgG (Sigma-Aldrich, S1265). Beads were then washed four times with 1 ml of ‘mES wash buffer’, before elution of the bound proteins at 95 °C for 3 minutes in 40 µl 1X LDS sample loading buffer.

For mass spectrometry analysis, 12-15 dishes were used per sample. As above, cells were resuspended in one volume of ‘mES lysis buffer’ containing protease inhibitors and 5 μM Propargylated-Ubiquitin, before digestion of chromosomal DNA and centrifugation. Subsequently, the clarified extract was divided into two 1.5 ml microfuge tubes, to each of which was added 30 μl slurry of M-270 epoxy beads coupled to rabbit IgG (prepared without addition of BSA). The beads were then washed four times with 1 ml of ‘mES wash buffer’, before the bound proteins were eluted at 95 °C for 3 minutes with 15 µl 1X LDS sample loading buffer. The eluate from the two tubes was then pooled together before loading on a 4–12% Bis-Tris NuPAGE gel (Life Technologies, NP0301), using 1× MOPS buffer. Gels were stained with colloidal Coomassie (ThermoFisher Scientific, Simply Blue, LC6060) and each sample lane was then cut into 40 bands before digestion with trypsin. Peptides were analysed by nano liquid chromatography tandem mass spectrometry with Thermofisher Orbitrap platforms (MS Bioworks, https://www.msbioworks.com/). Data were analysed using Scaffold software (Proteome Software Inc, USA).

### Immunofluorescence and spinning disk confocal microscopy

Mouse ES cells were grown on chambered 4-well slides (µ-slide 4 well, 80426, Ibidi) before fixation in 4 % (w/v) paraformaldehyde (ThermoFisher Scientific, J19943.K2) in PBS for 10 minutes at room temperature. Fixed cells were then washed with PBS supplemented with 3 % (w/v) BSA, 1 mM MgCl_2_ and 1 mM CaCl_2_ for 30 minutes.

Chromosomal DNA was stained by incubation overnight at 4°C in a mixture of 1µg / ml Hoechst 333423 (Sigma, B2261), 3 %(w/v) BSA, 1 mM MgCl_2_ and 1 mM CaCl_2_ in PBS, before washing three times with PBS containing 3 % BSA, 1 mM MgCl_2_ and 1 mM CaCl_2_.

For immunofluorescence analysis, the fixed and washed cells were permeabilised by detergent extraction for 10 minutes in PBS supplemented with 3 % (w/v) BSA, 1 mM MgCl_2_, 1 mM CaCl_2_ and 0.5 % (v/v) Triton X-100. The cells were then incubated for 30 minutes with a blocking mixture that comprised PBS plus 3 % (w/v) BSA, 1 mM MgCl_2_ and 1 mM CaCl_2_. After, cells were then incubated at room temperature for 1 hour in a 1:500 dilution of antibody specific for CTF4 or CLASPIN, in PBS supplemented with 3 % (w/v) BSA, 1 mM MgCl_2_ and 1 mM CaCl_2_. Samples were then washed three times with 3 % (w/v) BSA, 1 mM MgCl_2_ and 1 mM CaCl_2_, before incubation for one hour at room temperature in the dark with a 1:2,000 dilution of secondary antibody (anti-sheep IgG coupled with Alexa Fluor^TM^ 594, Invitrogen, A-11016), in PBS supplemented with 3 % (w/v) BSA, 1 mM MgCl_2_, 1 mM CaCl_2_ and 1 µg / ml Hoechst 33342. Finally, the cells were washed three times with PBS containing 1 mM MgCl_2_ and 1 mM CaCl_2_, before mounting with 700 µl of PBS containing 1 mM MgCl_2_ and 1 mM CaCl_2_.

Confocal images were acquired with a Zeiss Cell Observer SD microscope and Yokogawa CSU-X1 spinning disk, using a HAMAMATSU C13440 camera with a PECON incubator, a 60X 1.4-NA Plan-Apochromat oil immersion objective, and the appropriate excitation and emission filter sets. Images were acquired using ‘ZEN blue’ software (Zeiss) and processed with ImageJ software (National Institutes of Health) as previously described ^66^.

### Yeast Two-Hybrid Assays

Budding yeast cells of the strain PJ69-4A (Table S5) were transformed with derivatives of the pGADT7 plasmid (fusion of protein of interest to the Gal4 activation domain; *LEU2* marker gene) and the pGBKT7 plasmid (fusion of the protein of interest to the Gal4 DNA binding domain; *TRP1* marker gene). For each experiment, five independent transformed colonies were resuspended together in water, diluted to 3.33 x 10^6^ cells / ml, and diluted again to 3.33 x 10^5^, 3.33 x 10^4^, and 3.33 x 10^3^ cells / ml. In each case, a 15 µl aliquot was plated in a series of spots that corresponded to 50,000, 5000, 500 and 50 cells. Plates contained 1.7%(w/v) agar, 2%(w/v) Glucose, 1x ‘Complete supplement drop-out’ -Ade-His-Leu-Met-Trp-Ura (FORMEDIUM, DCS1511), 1x ‘BD Difco™ Yeast Nitrogen Base without Amino Acids and Ammonium Sulfate’ (BD, 233520), supplemented where necessary by 0.04 mg/ml adenine and/or 0.04 mg/ml histidine. Plates were incubated at 30 °C for 2-3 days as indicated, before imaging with a scanner.

### *In silico* modelling of protein structure and interactions using AlphaFold

Structural predictions were performed using AlphaFold-Multimer ^41^ and AlphaFold 3 ^67^, via ColabFold ^68^ and the AlphaFold 3 server (https://golgi.sandbox.google.com/about). Predicted structures were then analysed with Chimera X (https://www.rbvi.ucsf.edu/chimerax/).

The AlphaFold 3 Analysis Tool (https://predictomes.org/tools/af3/) was used to generate Predicted Alignment Error (PAE) plots of AlphaFold 3 prediction data.

### Quantification and Statistical Analysis

GraphPad Prism (GraphPad Software) was used to perform statistical analysis.

---

**Figure S1.**
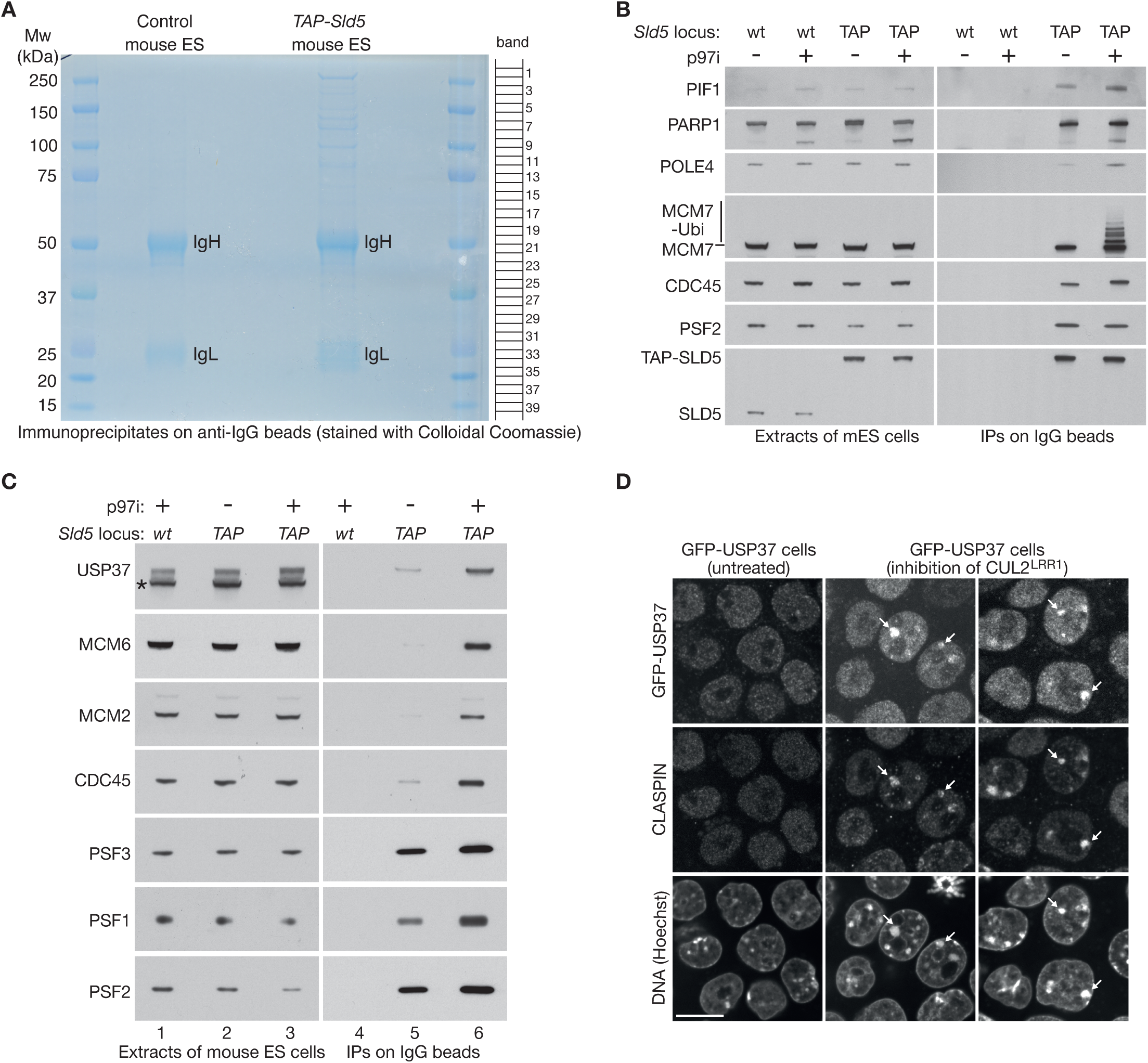
Proteins associated with the CMG helicase in mouse ES cells were analysed by mass spectrometry. (**A**) Extracts of control and TAP-Sld5 cells were incubated with magnetic beads coated with IgG. Bound proteins were eluted and resolved in a 4-12% gradient gel, which was stained with colloidal Coomassie blue. Each lane was then cut into 40 bands as shown before analysis by mass spectrometry. See Supplementary Tables S1-S4. (**B**-**C**) To validate the results of the mass spectrometry analysis, mouse ES cells of the indicated genotypes were treated as shown with 5 µM CB-5083 (p97i) to inhibit the p97 ATPase, before incubation of cell extracts with magnetic beads coated with IgG. The indicated factors were monitored by immunoblotting. (**D**) *GFP-Usp37* mouse ES cells were processed as for Figure 1B. The arrows indicate the accumulation of CLASPIN and GFP-USP37 on patches of constitutive heterochromatin, in response to inhibition of CUL2^LRR1^. The scale bar corresponds to 10 µm.

**Figure S2.**
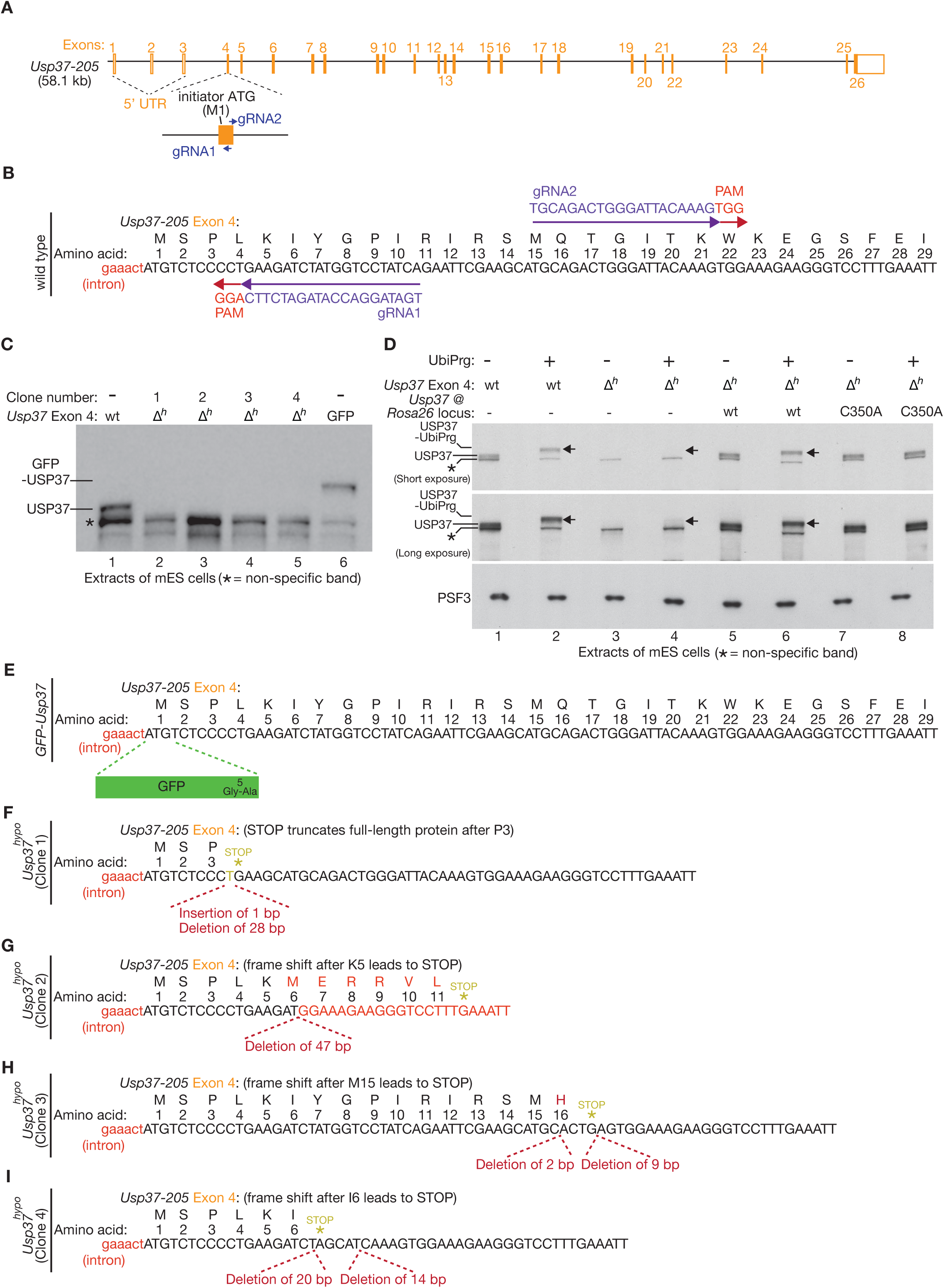
Using CRISPR-Cas9 to generate the *Usp37^hypo^* and *GFP-Usp37* alleles in mouse ES cells. (**A**) Illustration of the genomic locus of the mouse *Usp37-205* isoform, which encodes the orthologue of human USP37. The location is indicated of two guide RNAs (gRNAs) that were used to target the Cas9-D10A nickase to the first coding exon (Exon 4). (**B**) DNA sequence of the start of the *Usp37-205* coding sequence in Exon 4, illustrating the sequence of the two gRNAs that were used to target the Cas9-D10A nickase to the locus. PAM = Protomer Adjacent Motif (of form ‘NGG’), which is present in the genomic sequence immediately after the 20-mer targeting sequence of each gRNA, and is required for Cas9 to cut the DNA template. (**C**) Immunoblot analysis of wild type mouse ES cells, cells with GFP inserted in front of the initiator codon of Usp37 in Exon 4 (*GFP-Usp37*), and the indicated clones of cells with small deletions around the initiator codon of *Usp37* in Exon 4 (‘*Δh’* = *Usp37^hypo^*). (**D**) The activity of USP37 in extracts of mouse ES cells was monitored by addition of Ubi-Prg, followed by SDS-PAGE and immunoblotting. Covalent attachment of Ubi-Prg to USP37 was revealed as a mobility shift (compare lane 2 with lane 1). CRISPR deletions at the start of the *Usp37* coding sequence impaired the expression of full-length USP37 (lanes 3-4, short exposure; ‘*Δh*’ = *Usp37^hypo^*). However, longer exposures revealed a small population of truncated USP37 in *Usp37^hypo^* cells, which was active as it reacted with Ubi-Prg (lanes 3-4, long exposure; shifted forms marked by arrow). *Usp37^hypo^* could be rescued by expression of wild type USP37 from the *Rosa26* locus (lanes 5-6, arrow marks shift + Ubi-Prg). In contrast, USP37-C350A could be expressed in *Usp37^hypo^* cells to a similar level to the wild type protein but was inactive (lanes 7-8, no shift was observed + Ubi-Prg), consistent with the data in Figure 3C-D. (**E**) DNA sequence of the start of the *Usp37* coding sequence in Usp37-205 Exon 4, illustrating the site at which GFP was inserted by CRISPR-Cas9, to create the *GFP-Usp37* allele. (**F**-**I**) DNA sequence analysis of *Usp37^hypo^* clones 1-4, produced by transfecting cells with gRNAs 1+2 from (B). The panels show the predicted consequences in each clone for the expression of full-length USP37 protein.

**Figure S3.**
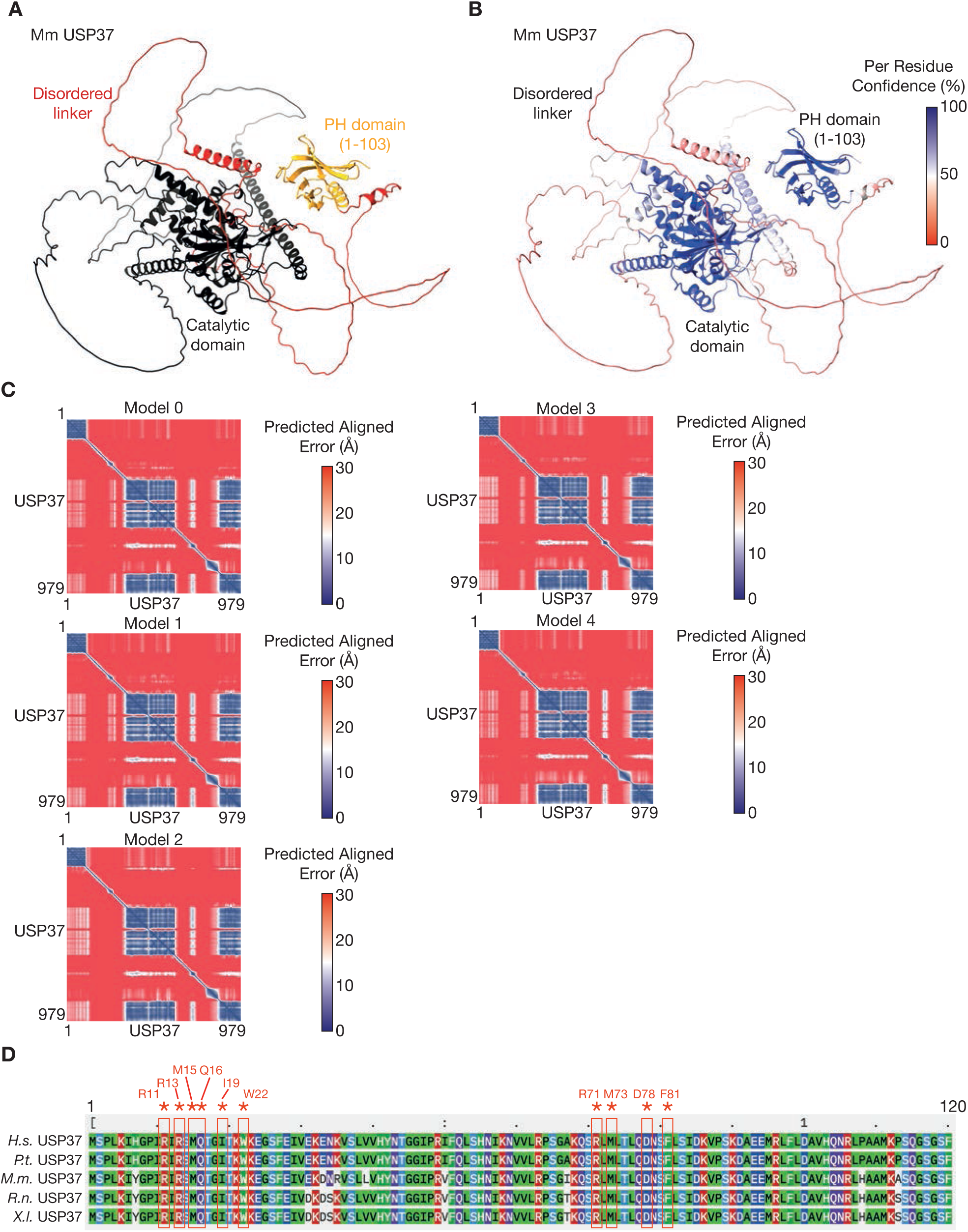
The PH domain of USP37 is separated from the catalytic domain by a long disordered linker. (**A**) AlphaFold 3 prediction of the structure of mouse USP37, showing the amino-terminal PH domain (1-103) in gold. Shown in red is the disordered linker (residues 104-349) that connects the PH domain to the catalytic domain (black). (**B**) The predicted model of USP37 from (A) is coloured according to Per Residue Confidence. (**C**) Predicted Alignment Error (PAE) plots for the five predictions of the USP37 structure by Alphafold3. (**D**) Sequence alignment for the amino terminus of USP37 containing the PH domain. The 10 conserved residues mutated in the USP37-10A allele are shown in red. *H.s.* = *Homo sapiens*; *P.t.* = *Pan troglodytes*; *M.m.* = *Mus musculus*; *R.n.* = *Rattus norvegicus*; *X.l.* = *Xenopus laevis*.

**Figure S4.**
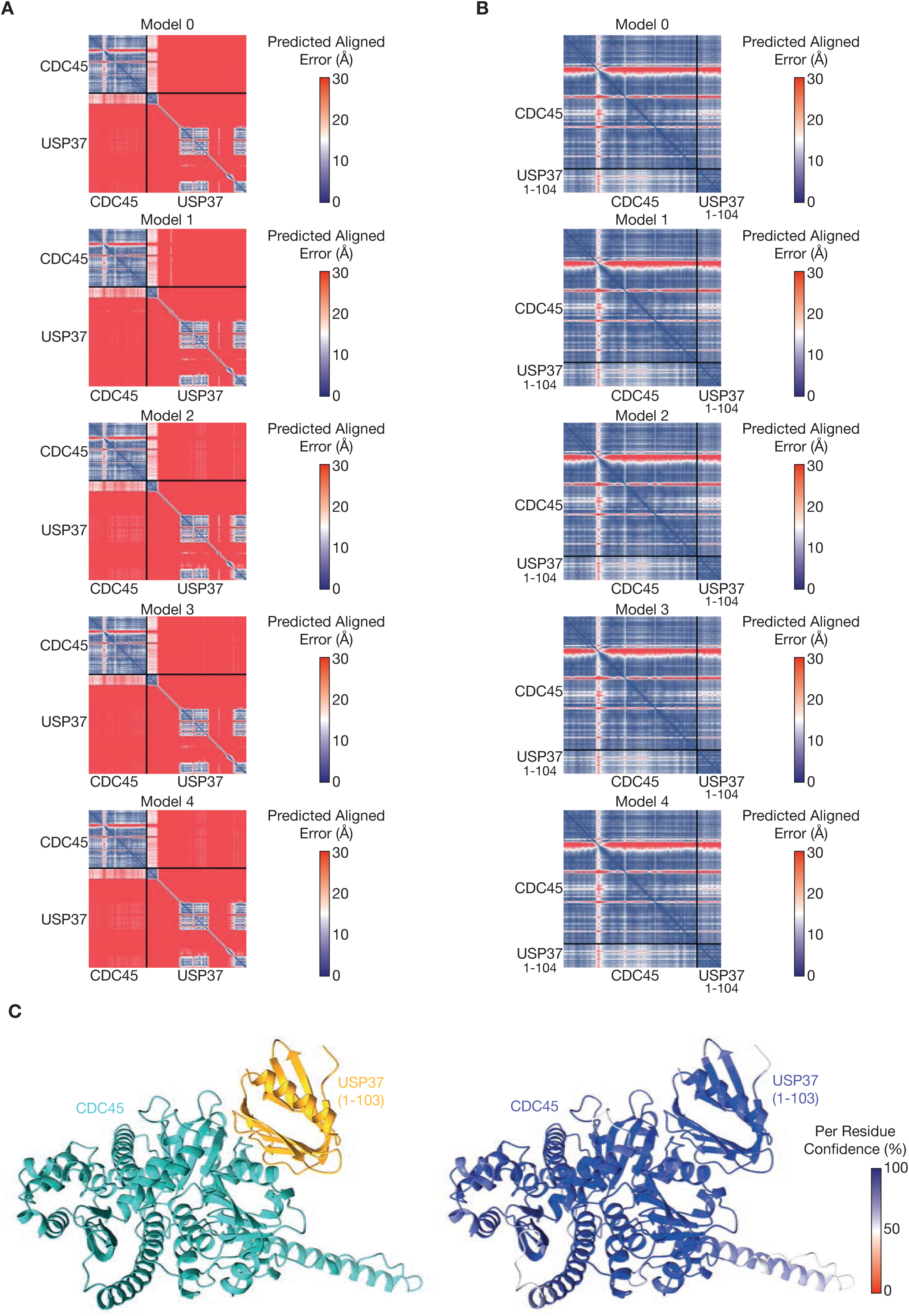
AlphaFold 3 prediction for binding of the USP37-PH domain to CDC45. (**A**) Predicted Alignment Error (PAE) plots for the five Alphafold3 predictions of the complex between full-length USP37 and CDC45. (**B**) Equivalent Predicted Alignment Error (PAE) plots for the five Alphafold3 predictions of the complex between the USP37-PH domain (1-103) and CDC45. (**B**) The predicted complex between CDC45 and the USP37-PH domain (left) is shown with colouring according to Per Residue Confidence (right).

**Figure S5.**
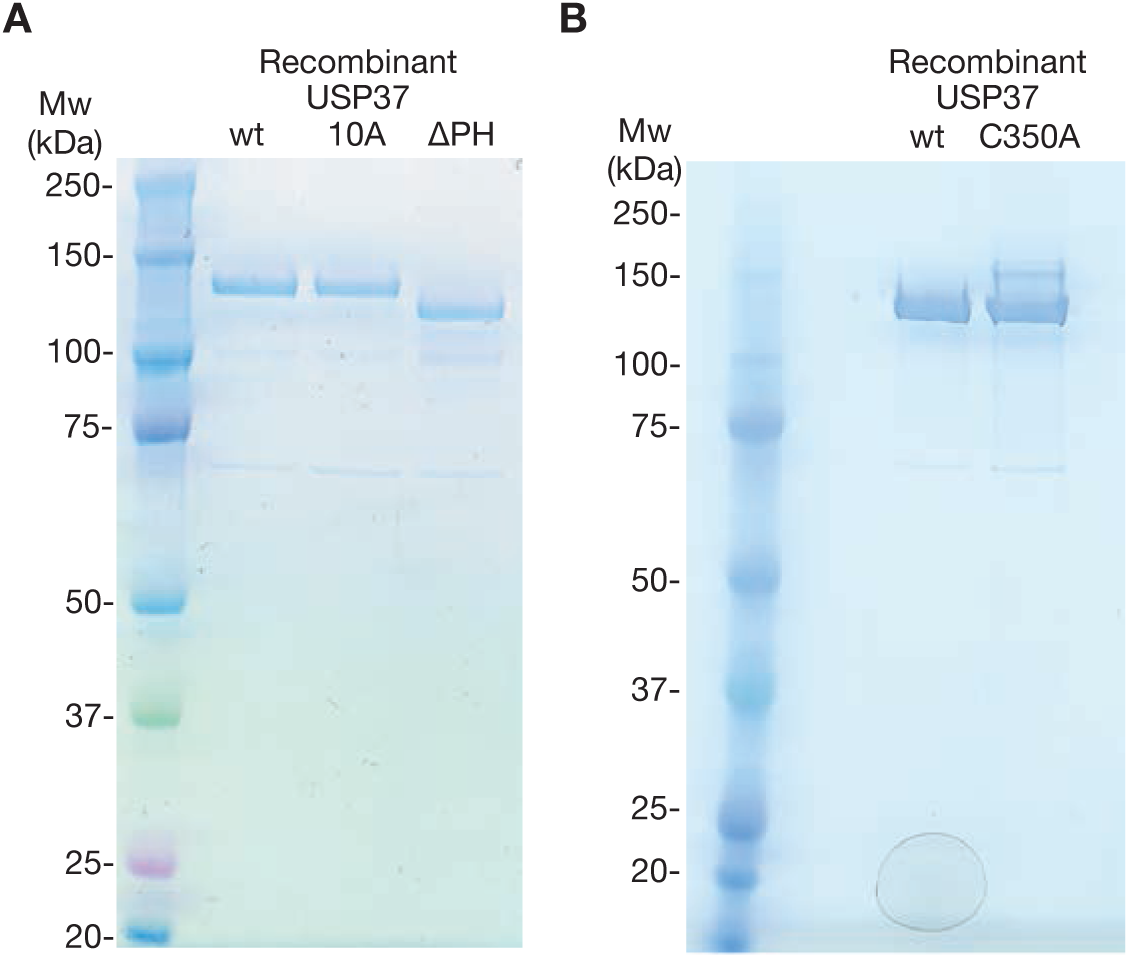
Recombinant mouse USP37 proteins after purification from budding yeast. (**A**) Mouse USP37, USP37-10A and USP37-ΔPH were expressed in budding yeast cells, purified and then analysed by SDS-PAGE, before staining with colloidal Coomassie blue. (**B**) Comparison of purified wild type USP37 and USP37-C350A. The upper band for USP37-C350A is likely to represent ubiquitylation of catalytically dead USP37.

**Figure S6.**
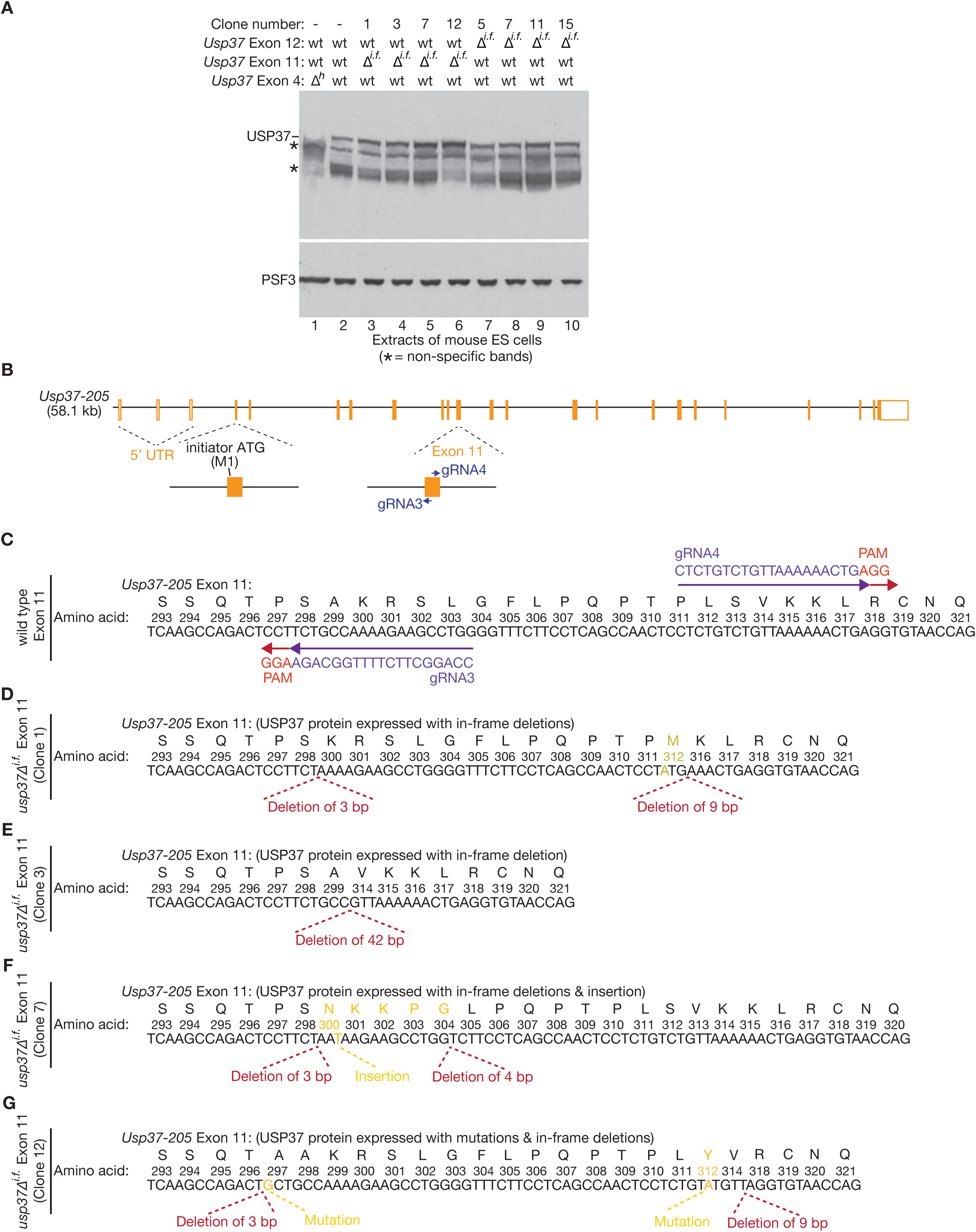
Targeting CRISPR-Cas9-D10A to *Usp37* Exon 11 leads to selection of viable clones that express USP37 with small in-frame deletions. (**A**) Immunoblot analysis of *Usp37^hypo^* (Exon 4 ‘Δh’, lane 1), wild type (lane 2) and four clones each of viable clones with CRISPR-Cas9 deletions in Exon 11 (lanes 3-6) or Exon 12 (lanes 7-10) or *Usp37*. ‘*Δ^i.f.^*’ = in-frame deletions in Exons 11 or 12 of *Usp37*. (**B**) Illustration of the *Usp37-205* locus in mouse ES cells. The location is indicated of two guide RNAs (gRNAs) that were used to create small deletions or insertions in Exon 11 (within the coding sequence for the catalytic domain), via targeting of the Cas9-D10A nickase and subsequent repair. (**C**) DNA sequence of the relevant region of *Usp37-205* Exon 11, illustrating the sequence of the two gRNAs that were used to target the Cas9-D10A nickase to the locus. PAM = Protomer Adjacent Motif (of form ‘NGG’), which is present in the genomic sequence immediately after the 20-mer targeting sequence of each gRNA and is required for Cas9 to cut the DNA template. (**D**-**G**) DNA sequence analysis of four viable clones with in-frame small deletions or mutations in Exon 11, produced by transfecting cells with gRNAs 3+4 from (B). The panels show the consequences in each clone for the expression of USP37 protein.

**Figure S7.**
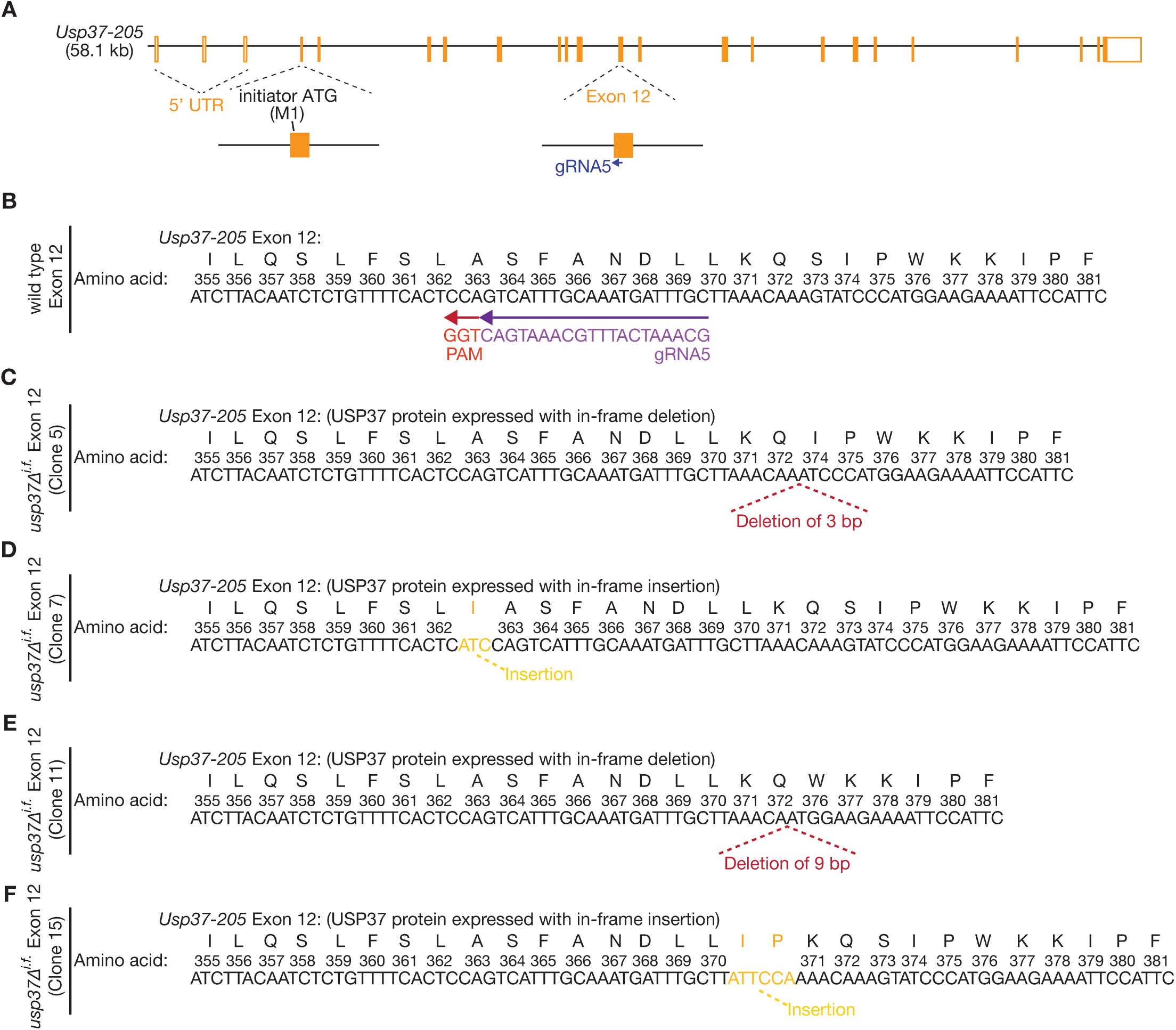
Targeting CRISPR-Cas9-D10A to *Usp37* Exon 12 leads to selection of viable clones that express USP37 with small in-frame deletions. (**A**) Illustration of the *Usp37* locus in mouse ES cells. The location is indicated of a single guide RNA (gRNA) that was used to create small deletions or insertions in Exon 12 (within the coding sequence for the catalytic domain), via targeting of wild type Cas9 and subsequent repair. (**B**) DNA sequence of the relevant region of *Usp37-205* Exon 12, illustrating the sequence of the gRNA that was used to target wild type Cas9 to the locus. PAM = Protomer Adjacent Motif (of form ‘NGG’), which is present in the genomic sequence immediately after the 20-mer targeting sequence of each gRNA and is required for Cas9 to cut the DNA template. (**C**-**F**) DNA sequence analysis of four viable clones with in-frame small deletions or mutations in Exon 12, produced by transfecting cells with gRNA5 from (B). The panels show the consequences in each clone for the expression of USP37 protein. ‘*Δ^i.f.^*’ = in-frame deletions in Exons 11 or 12 of *Usp37-205*.

**Figure S8.**
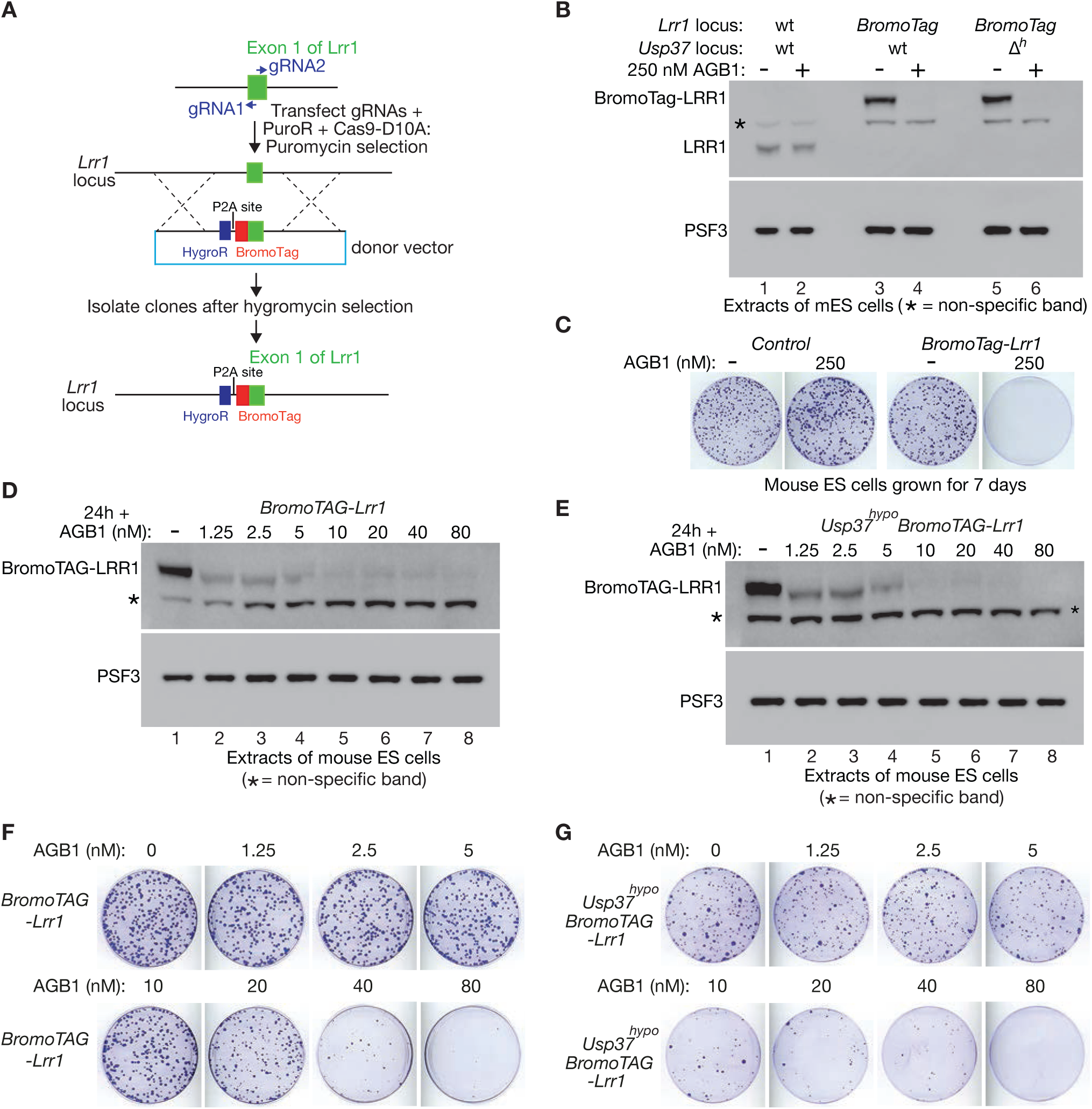
Fusion to BromoTag allows titratable depletion of the essential LRR1 protein in mouse ES cells. (**A**) Scheme for insertion of the BromoTag before the initiator codon in Exon 1 of *Lrr1* in mouse ES cells. (**B**) Immunoblot analysis of cells of the indicated genotypes, treated as shown for 3 hours with 250 nM AGB1. ‘*Δh*’ = *Usp37^hypo^*. (**C**) Cells were grown as shown in the presence or absence of AGB1, before staining of colonies with crystal violet. (**D**-**E**) Cells of the indicated genotypes were grown for 24 h in the presence of the indicated concentrations of AGB1, before immunoblot analysis to monitor the level of BromoTag-LRR1. (**F**-**G**) Cells were grown for seven days as shown in the presence of the indicated concentrations of AGB1, before staining of colonies with crystal violet.

**Table S1.** Mass spectrometry data showing that the USP37 deubiquitylase copurifies with the CMG helicase (and associated factors) from extracts of mouse embryonic stem cells. This is the first of four independent experiments. The others are in Tables S2-S4.

| **Protein** | **Complex** | **Spectral counts (control)** | **Spectral counts (TAP-SLD5)** |
| --- | --- | --- | --- |
| **Replisome components** |  |  |  |
| PSF1 (23kDa) | CMG (GINS) | 12 | 1174 |
| PSF2 (21kDa) | CMG (GINS) | 2 | 605 |
| PSF3 (22kDa) | CMG (GINS) | 0 | 744 |
| SLD5 (26kDa) | CMG (GINS) | 0 | 1348 |
| CDC45 (65kDa) | CMG | 0 | 394 |
| MCM2 (102kDa) | CMG (MCM2-7) | 130 | 1311 |
| MCM3 (92kDa) | CMG (MCM2-7) | 97 | 996 |
| MCM4 (97kDa) | CMG (MCM2-7) | 10 | 898 |
| MCM5 (82kDa) | CMG (MCM2-7) | 55 | 780 |
| MCM6 (93kDa) | CMG (MCM2-7) | 13 | 718 |
| MCM7 (81kDa) | CMG (MCM2-7) | 50 | 503 |
| CTF4 (124kDa) |  | 5 | 1026 |
| TIMELESS (138kDa) | TIMELESS-TIPIN | 2 | 959 |
| TIPIN (31kDa) | TIMELESS-TIPIN | 0 | 125 |
| CLASPIN (147kDa) |  | 0 | 926 |
| SUPT16H (120kDa) | FACT | 0 | 664 |
| SSRP1 (81kDa) | FACT | 0 | 375 |
| POLE1 (262kDa) | Pol $\epsilon$ | 12 | 1185 |
| POLE2<br>(59kDa) | Pol $\epsilon$ | 0 | 186 |
| POLE3<br>(17kDa) | Pol $\epsilon$ | 0 | 22 |
| POLE4<br>(12kDa) | Pol $\epsilon$ | 0 | 25 |
| POLA1<br>(167kDa) | Pol $\alpha$ | 7 | 663 |
| POLA2<br>(66kDa) | Pol $\alpha$ | 0 | 103 |
| PRIM1<br>(48kDa) | Pol $\alpha$ | 0 | 195 |
| PRIM2<br>(58kDa) | Pol $\alpha$ | 0 | 238 |
| CHTF18<br>(108kDa) | CTF18-RFC | 0 | 760 |
| CHTF8<br>(13kDa) | CTF18-RFC | 0 | 40 |
| DSCC1<br>(46kDa) | CTF18-RFC | 0 | 104 |
| RFC2<br>(39kDa) | CTF18-RFC | 14 | 121 |
| RFC3<br>(41kDa) | CTF18-RFC | 21 | 205 |
| RFC4<br>(40kDa) | CTF18-RFC | 23 | 159 |
| RFC5<br>(38kDa) | CTF18-RFC | 9 | 111 |
| **Ubiquitin**<br>**biology** |  |  |  |
| USP37<br>(110kDa) |  | 0 | 101 |
| TRAIP<br>(53kDa) |  | 0 | 47 |
| **Selected**<br>**other**<br>**factors** |  |  |  |
| PARP1<br>(111kDa) |  | 68 | 618 |
| RIF1<br>(266kDa) |  | 15 | 264 |
| PIF1<br>(71kDa) |  | 0 | 177 |
| RTEL1<br>(134kDa) |  | 0 | 128 |
| TONSL<br>(151kDa) | TONSL-MMS22L | 3 | 73 |
| MMS22L<br>(141kDa) | TONSL-MMS22L | 2 | 39 |
| SENP6<br>(127kDa) |  | 0 | 72 |
| EXO1<br>(92kDa) |  | 0 | 36 |
| **DNA**<br>**replication**<br>**initiation**<br>**factors** |  |  |  |
| TRESLIN<br>(208kDa) |  | 0 | 135 |
| MCM10<br>(98kDa) |  | 0 | 131 |
| MTBP<br>(92kDa) |  | 0 | 48 |
| DONSON<br>(62kDa) |  | 0 | 36 |
| TOPBP1<br>(169kDa) |  | 0 | 36 |

**Table S2.** Mass spectrometry data showing that the USP37 deubiquitylase copurifies with the CMG helicase (and associated factors) from extracts of mouse embryonic stem cells. This is the first of four independent experiments. The others are in Tables S1 and S3-S4.

| **Protein** | **Complex** | **Spectral counts (control)** | **Spectral counts (TAP-SLD5)** |
| --- | --- | --- | --- |
| **Replisome components** |  |  |  |
| PSF1 (23kDa) | CMG (GINS) | 25 | 1192 |
| PSF2 (21kDa) | CMG (GINS) | 13 | 530 |
| PSF3 (22kDa) | CMG (GINS) | 60 | 666 |
| SLD5 (26kDa) | CMG (GINS) | 6 | 1499 |
| CDC45 (65kDa) | CMG | 0 | 335 |
| MCM2 (102kDa) | CMG (MCM2-7) | 103 | 1644 |
| MCM3 (92kDa) | CMG (MCM2-7) | 74 | 1230 |
| MCM4 (97kDa) | CMG (MCM2-7) | 41 | 1048 |
| MCM5 (82kDa) | CMG (MCM2-7) | 59 | 837 |
| MCM6 (93kDa) | CMG (MCM2-7) | 59 | 844 |
| MCM7 (81kDa) | CMG (MCM2-7) | 56 | 692 |
| CTF4 (124kDa) |  | 14 | 993 |
| TIMELESS (138kDa) | TIMELESS-TIPIN | 2 | 813 |
| TIPIN (31kDa) | TIMELESS-TIPIN | 0 | 130 |
| CLASPIN (147kDa) |  | 0 | 1327 |
| SUPT16H (120kDa) | FACT | 0 | 614 |
| SSRP1 (81kDa) | FACT | 0 | 386 |
| POLE1 (262kDa) | Pol $\epsilon$ | 15 | 1420 |
| POLE2<br>(59kDa) | Pol $\epsilon$ | 0 | 244 |
| POLE3<br>(17kDa) | Pol $\epsilon$ | 0 | 64 |
| POLE4<br>(12kDa) | Pol $\epsilon$ | 0 | 49 |
| POLA1<br>(167kDa) | Pol $\alpha$ | 3 | 738 |
| POLA2<br>(66kDa) | Pol $\alpha$ | 4 | 128 |
| PRIM1<br>(48kDa) | Pol $\alpha$ | 0 | 162 |
| PRIM2<br>(58kDa) | Pol $\alpha$ | 0 | 237 |
| CHTF18<br>(108kDa) | CTF18-RFC | 0 | 705 |
| CHTF8<br>(13kDa) | CTF18-RFC | 0 | 18 |
| DSCC1<br>(46kDa) | CTF18-RFC | 0 | 134 |
| RFC2<br>(39kDa) | CTF18-RFC | 18 | 217 |
| RFC3<br>(41kDa) | CTF18-RFC | 24 | 245 |
| RFC4<br>(40kDa) | CTF18-RFC | 12 | 195 |
| RFC5<br>(38kDa) | CTF18-RFC | 12 | 143 |
| **Ubiquitin**<br>**biology** |  |  |  |
| USP37<br>(110kDa) |  | 0 | 64 |
| TRAIP<br>(53kDa) |  | 0 | 39 |
| **Selected**<br>**other**<br>**factors** |  |  |  |
| PARP1<br>(111kDa) |  | 56 | 720 |
| RIF1<br>(266kDa) |  | 9 | 295 |
| PIF1<br>(71kDa) |  | 0 | 239 |
| RTEL1<br>(134kDa) |  | 0 | 81 |
| TONSL<br>(151kDa) | TONSL-MMS22L | 0 | 71 |
| MMS22L<br>(141kDa) | TONSL-MMS22L | 6 | 36 |
| SENP6<br>(127kDa) |  | 0 | 78 |
| EXO1<br>(92kDa) |  | 0 | 48 |
| **DNA**<br>**replication**<br>**initiation**<br>**factors** |  |  |  |
| TRESLIN<br>(208kDa) |  | 0 | 212 |
| MCM10<br>(98kDa) |  | 0 | 72 |
| MTBP<br>(92kDa) |  | 0 | 70 |
| DONSON<br>(62kDa) |  | 0 | 15 |
| TOPBP1<br>(169kDa) |  | 4 | 44 |

**Table S3.** Mass spectrometry data showing that the USP37 deubiquitylase copurifies with the CMG helicase (and associated factors) from extracts of mouse embryonic stem cells. This is the third of four independent experiments. The others are in Tables S1-S2 and S4.

| **Protein** | **Complex** | **Spectral counts (control)** | **Spectral counts (*TAP-Sld5*)** | **Spectral counts (*TAP-Sld5* + p97i)** |
| --- | --- | --- | --- | --- |
| **Replisome components** |  |  |  |  |
| PSF1 (23kDa) | CMG (GINS) | 0 | 387 | 522 |
| PSF2 (21kDa) | CMG (GINS) | 0 | 231 | 323 |
| PSF3 (22kDa) | CMG (GINS) | 0 | 200 | 299 |
| SLD5 (26kDa) | CMG (GINS) | 0 | 536 | 793 |
| CDC45 (65kDa) | CMG | 0 | 337 | 782 |
| MCM2 (102kDa) | CMG (MCM2-7) | 0 | 903 | 1525 |
| MCM3 (92kDa) | CMG (MCM2-7) | 20 | 910 | 1597 |
| MCM4 (97kDa) | CMG (MCM2-7) | 0 | 626 | 1121 |
| MCM5 (82kDa) | CMG (MCM2-7) | 15 | 794 | 1493 |
| MCM6 (93kDa) | CMG (MCM2-7) | 5 | 841 | 1630 |
| MCM7 (81kDa) | CMG (MCM2-7) | 29 | 641 | 1545 |
| CTF4 (124kDa) |  | 6 | 1356 | 2298 |
| TIMELESS (138kDa) | TIMELESS-TIPIN | 0 | 783 | 1923 |
| TIPIN (31kDa) | TIMELESS-TIPIN | 0 | 87 | 272 |
| CLASPIN (147kDa) |  | 0 | 994 | 1764 |
| SUPT16H (120kDa) | FACT | 0 | 547 | 1512 |
| SSRP1 (81kDa) | FACT | 0 | 140 | 429 |
| POLE1 (262kDa) | Pol $\epsilon$ | 34 | 1842 | 2896 |
| POLE2<br>(59kDa) | Pol $\epsilon$ | 0 | 252 | 443 |
| POLE3<br>(17kDa) | Pol $\epsilon$ | 0 | 22 | 30 |
| POLE4<br>(12kDa) | Pol $\epsilon$ | 0 | 17 | 24 |
| POLA1<br>(167kDa) | Pol $\alpha$ | 0 | 636 | 1399 |
| POLA2<br>(66kDa) | Pol $\alpha$ | 0 | 85 | 235 |
| PRIM1<br>(48kDa) | Pol $\alpha$ | 0 | 148 | 382 |
| PRIM2<br>(58kDa) | Pol $\alpha$ | 0 | 154 | 466 |
| CTF18<br>(108kDa) | CTF18-RFC | 0 | 539 | 1044 |
| CTF8<br>(13kDa) | CTF18-RFC | 0 | 9 | 20 |
| DSCC1<br>(46kDa) | CTF18-RFC | 0 | 92 | 247 |
| RFC2<br>(39kDa) | CTF18-RFC | 2 | 165 | 300 |
| RFC3<br>(41kDa) | CTF18-RFC | 0 | 225 | 428 |
| RFC4<br>(40kDa) | CTF18-RFC | 0 | 167 | 376 |
| RFC5<br>(38kDa) | CTF18-RFC | 0 | 118 | 214 |
| **Ubiquitin**<br>**biology** |  |  |  |  |
| USP37<br>(110kDa) |  | 0 | 55 | 429 |
| TRAIP<br>(53kDa) |  | 0 | 2 | 26 |
| **Selected**<br>**other**<br>**factors** |  |  |  |  |
| PARP1<br>(111kDa) |  | 6 | 650 | 1267 |
| RIF1<br>(266kDa) |  | 0 | 39 | 373 |
| PIF1<br>(71kDa) |  | 0 | 150 | 179 |
| RTEL1<br>(134kDa) |  | 0 | 42 | 78 |
| TONSL<br>(151kDa) | TONSL-MMS22L | 0 | 70 | 300 |
| MMS22L<br>(141kDa) | TONSL-MMS22L | 0 | 55 | 170 |
| SENP6<br>(127kDa) |  | 0 | 16 | 49 |
| EXO1<br>(92kDa) |  | 0 | 37 | 52 |
| **DNA**<br>**replication**<br>**initiation**<br>**factors** |  |  |  |  |
| TRESLIN<br>(208kDa) |  | 0 | 262 | 718 |
| MCM10<br>(98kDa) |  | 0 | 33 | 248 |
| MTBP<br>(92kDa) |  | 0 | 93 | 237 |
| DONSON<br>(62kDa) |  | 0 | 12 | 23 |
| TOPBP1<br>(169kDa) |  | 0 | 41 | 29 |

**Table S4.** Mass spectrometry data showing that the USP37 deubiquitylase copurifies with the CMG helicase (and associated factors) from extracts of mouse embryonic stem cells. This is the last of four independent experiments. The others are in Tables S1-S3.

| **Protein** | **Complex** | **Spectral counts (control)** | **Spectral counts (control, +p97i)** | **Spectral counts (TAP-Sld5)** | **Spectral counts (TAP-Sld5 +p97i)** |
| --- | --- | --- | --- | --- | --- |
| **Replisome components** |  |  |  |  |  |
| PSF1 (23kDa) | CMG (GINS) | 0 | 21 | 565 | 584 |
| PSF2 (21kDa) | CMG (GINS) | 0 | 0 | 232 | 256 |
| PSF3 (22kDa) | CMG (GINS) | 11 | 44 | 489 | 443 |
| SLD5 (26kDa) | CMG (GINS) | 108 | 144 | 1701 | 1731 |
| CDC45 (65kDa) | CMG | 0 | 0 | 460 | 613 |
| MCM2 (102kDa) | CMG (MCM2-7) | 5 | 46 | 1470 | 1996 |
| MCM3 (92kDa) | CMG (MCM2-7) | 5 | 36 | 1446 | 2007 |
| MCM4 (97kDa) | CMG (MCM2-7) | 10 | 22 | 1070 | 1576 |
| MCM5 (82kDa) | CMG (MCM2-7) | 0 | 30 | 1174 | 1764 |
| MCM6 (93kDa) | CMG (MCM2-7) | 10 | 39 | 1168 | 1822 |
| MCM7 (81kDa) | CMG (MCM2-7) | 10 | 26 | 786 | 1617 |
| CTF4 (124kDa) |  | 0 | 2 | 1993 | 2553 |
| TIMELESS (138kDa) | TIMELESS-TIPIN | 0 | 0 | 1261 | 1899 |
| TIPIN (31kDa) | TIMELESS-TIPIN | 0 | 0 | 112 | 160 |
| CLASPIN (147kDa) |  | 0 | 0 | 1008 | 1194 |
| SUPT16H (120kDa) | FACT | 0 | 0 | 793 | 982 |
| SSRP1 (81kDa) | FACT | 0 | 0 | 333 | 568 |
| POLE1 (262kDa) | Pol $\epsilon$ | 0 | 0 | 2011 | 3093 |
| POLE2<br>(59kDa) | Pol $\epsilon$ | 0 | 0 | 290 | 442 |
| POLE3<br>(17kDa) | Pol $\epsilon$ | 0 | 0 | 27 | 26 |
| POLE4<br>(12kDa) | Pol $\epsilon$ | 0 | 0 | 30 | 44 |
| POLA1<br>(167kDa) | Pol $\alpha$ | 0 | 0 | 863 | 1690 |
| POLA2<br>(66kDa) | Pol $\alpha$ | 0 | 0 | 147 | 327 |
| PRIM1<br>(48kDa) | Pol $\alpha$ | 0 | 3 | 185 | 427 |
| PRIM2<br>(58kDa) | Pol $\alpha$ | 0 | 0 | 322 | 608 |
| CTF18<br>(108kDa) | CTF18-RFC | 0 | 0 | 1033 | 1288 |
| CTF8<br>(13kDa) | CTF18-RFC | 0 | 0 | 29 | 42 |
| DSCC1<br>(46kDa) | CTF18-RFC | 0 | 0 | 123 | 267 |
| RFC2<br>(39kDa) | CTF18-RFC | 8 | 13 | 260 | 440 |
| RFC3<br>(41kDa) | CTF18-RFC | 3 | 7 | 258 | 483 |
| RFC4<br>(40kDa) | CTF18-RFC | 7 | 8 | 207 | 355 |
| RFC5<br>(38kDa) | CTF18-RFC | 6 | 5 | 160 | 283 |
| **Ubiquitin**<br>**biology** |  |  |  |  |  |
| USP37<br>(110kDa) |  | 0 | 0 | 129 | 447 |
| TRAIP<br>(53kDa) |  | 0 | 0 | 4 | 23 |
| **Selected**<br>**other**<br>**factors** |  |  |  |  |  |
| PARP1<br>(111kDa) |  | 3 | 22 | 686 | 1237 |
| RIF1<br>(266kDa) |  | 11 | 37 | 158 | 283 |
| PIF1<br>(71kDa) |  | 0 | 0 | 120 | 132 |
| RTEL1<br>(134kDa) |  | 0 | 0 | 33 | 93 |
| TONSL<br>(151kDa) | TONSL-MMS22L | 0 | 0 | 73 | 244 |
| MMS22L<br>(141kDa) | TONSL-MMS22L | 0 | 0 | 51 | 143 |
| SENP6<br>(127kDa) |  | 0 | 0 | 23 | 56 |
| EXO1<br>(92kDa) |  | 0 | 0 | 33 | 79 |
| **DNA**<br>**replication**<br>**initiation**<br>**factors** |  |  |  |  |  |
| TRESLIN<br>(208kDa) |  | 0 | 0 | 250 | 630 |
| MCM10<br>(98kDa) |  | 0 | 0 | 26 | 273 |
| MTBP<br>(92kDa) |  | 0 | 0 | 111 | 284 |
| DONSON<br>(62kDa) |  | 0 | 0 | 8 | 11 |
| TOPBP1<br>(169kDa) |  | 0 | 0 | 42 | 21 |

**Table S5.**
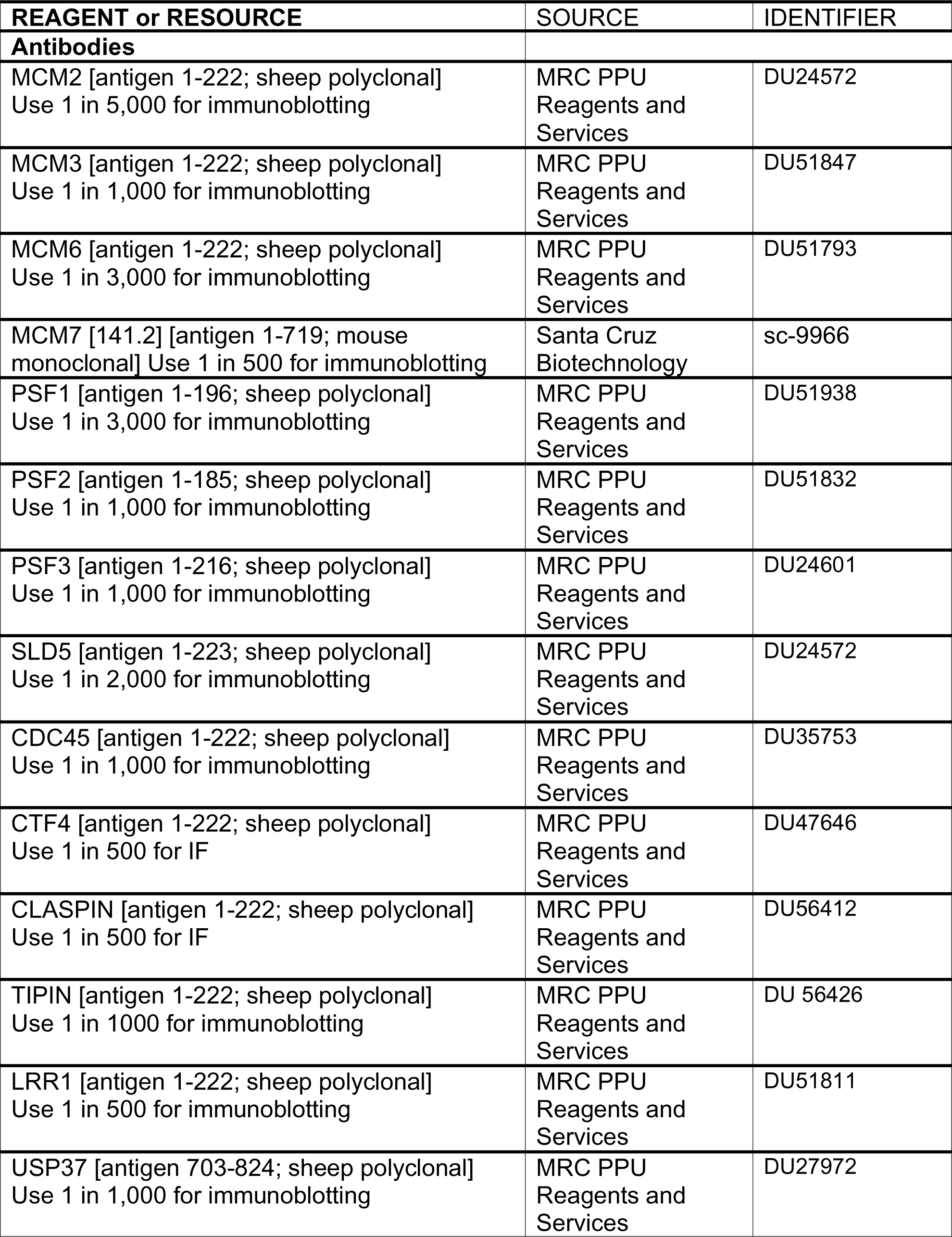
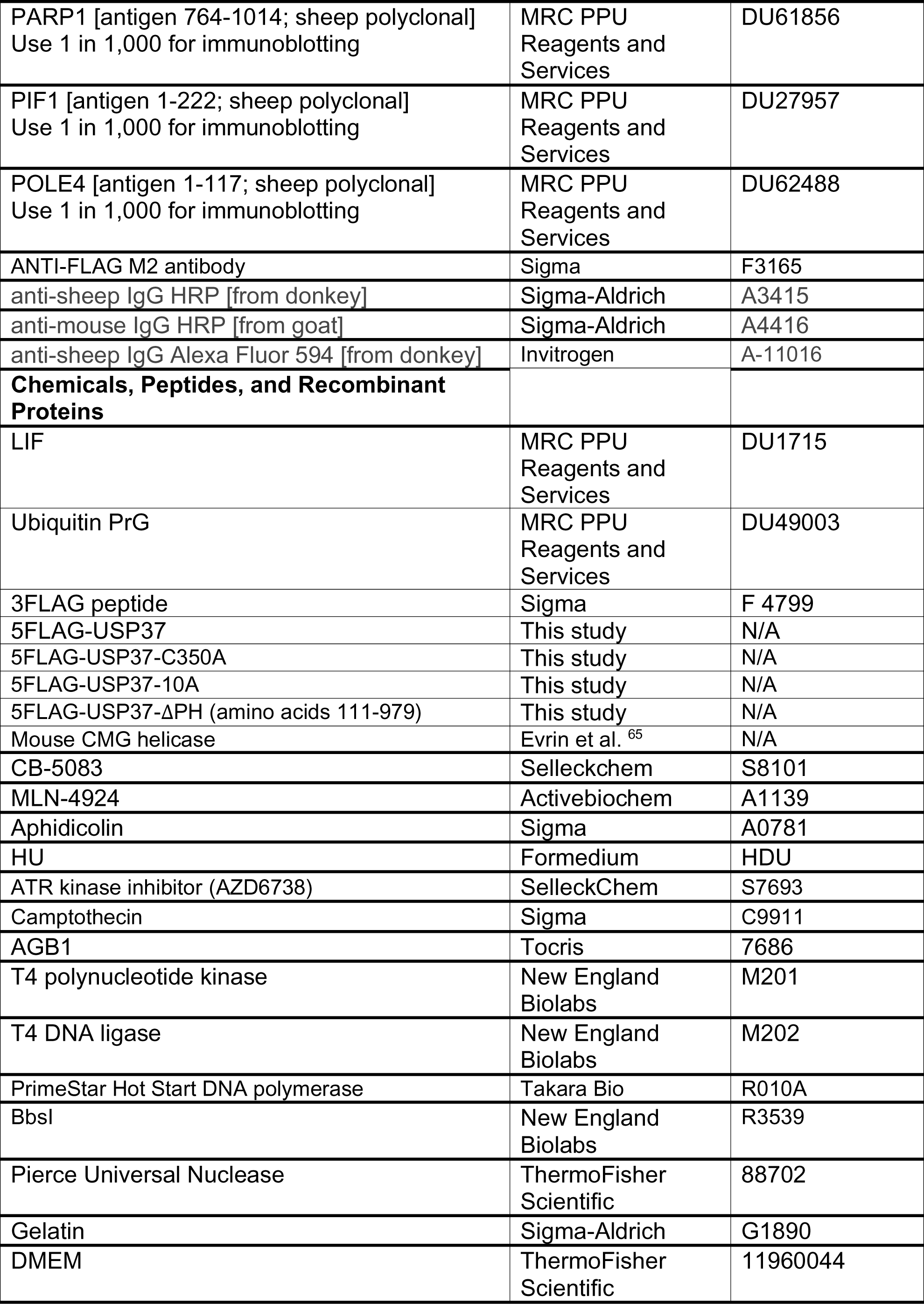
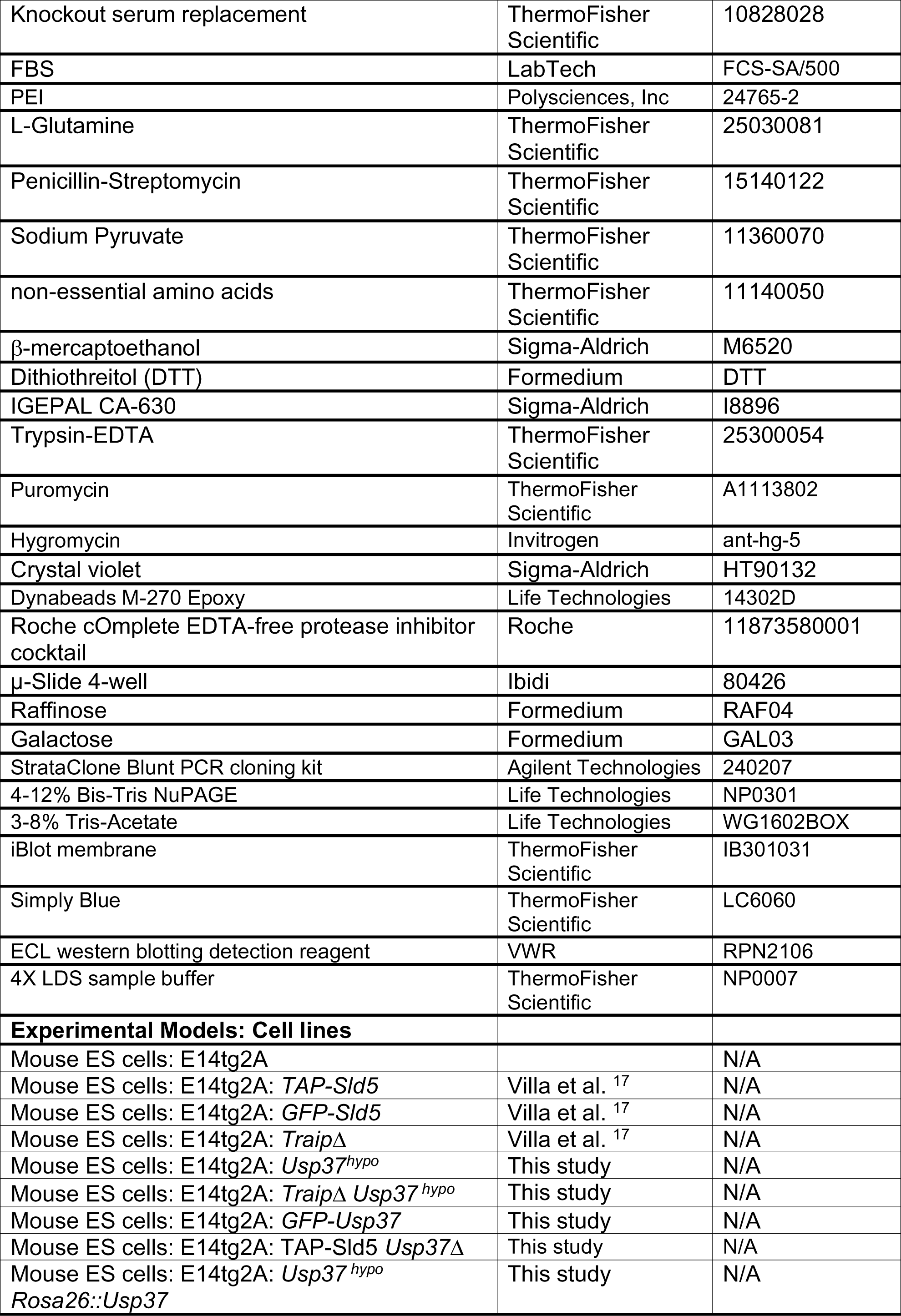
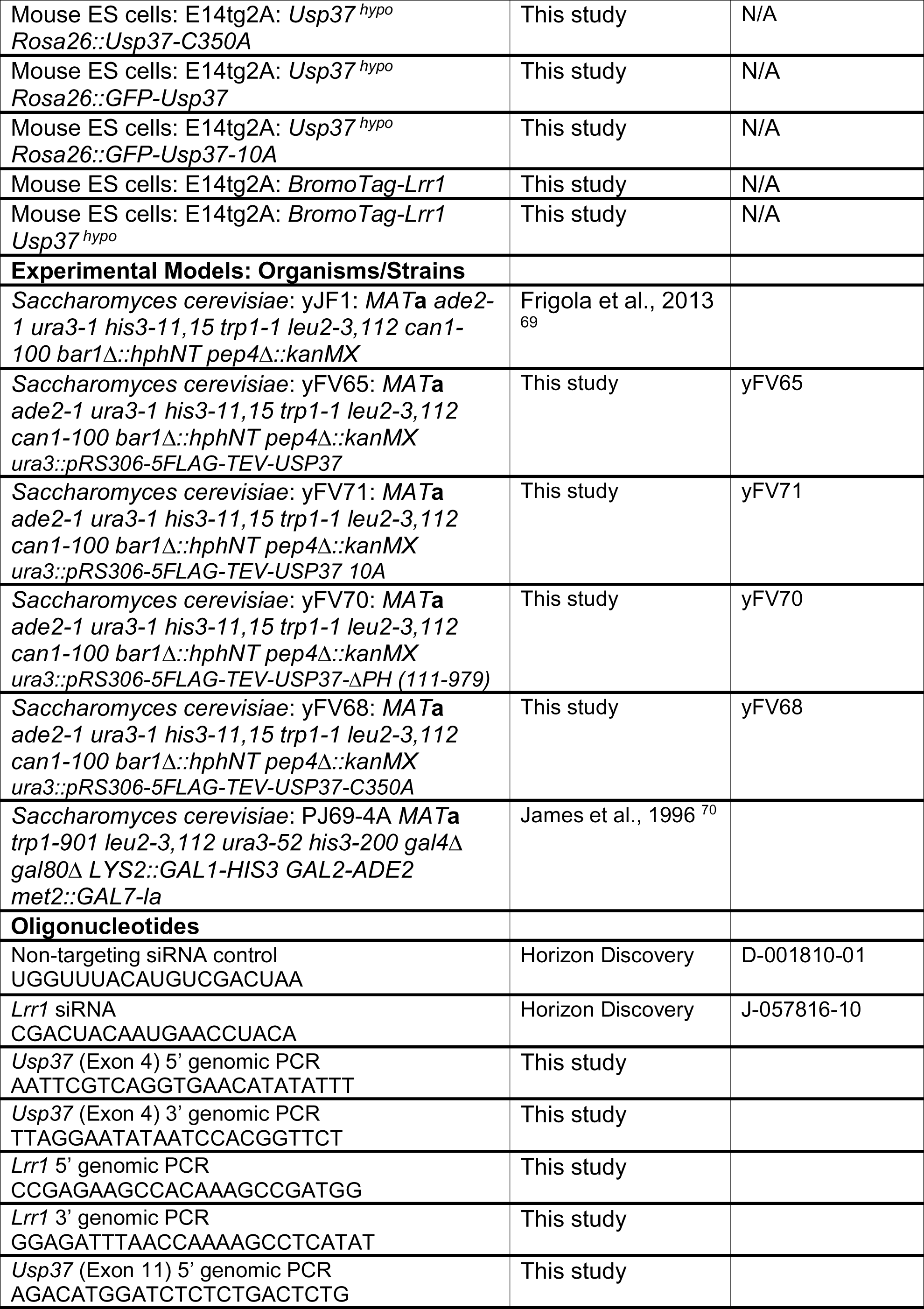
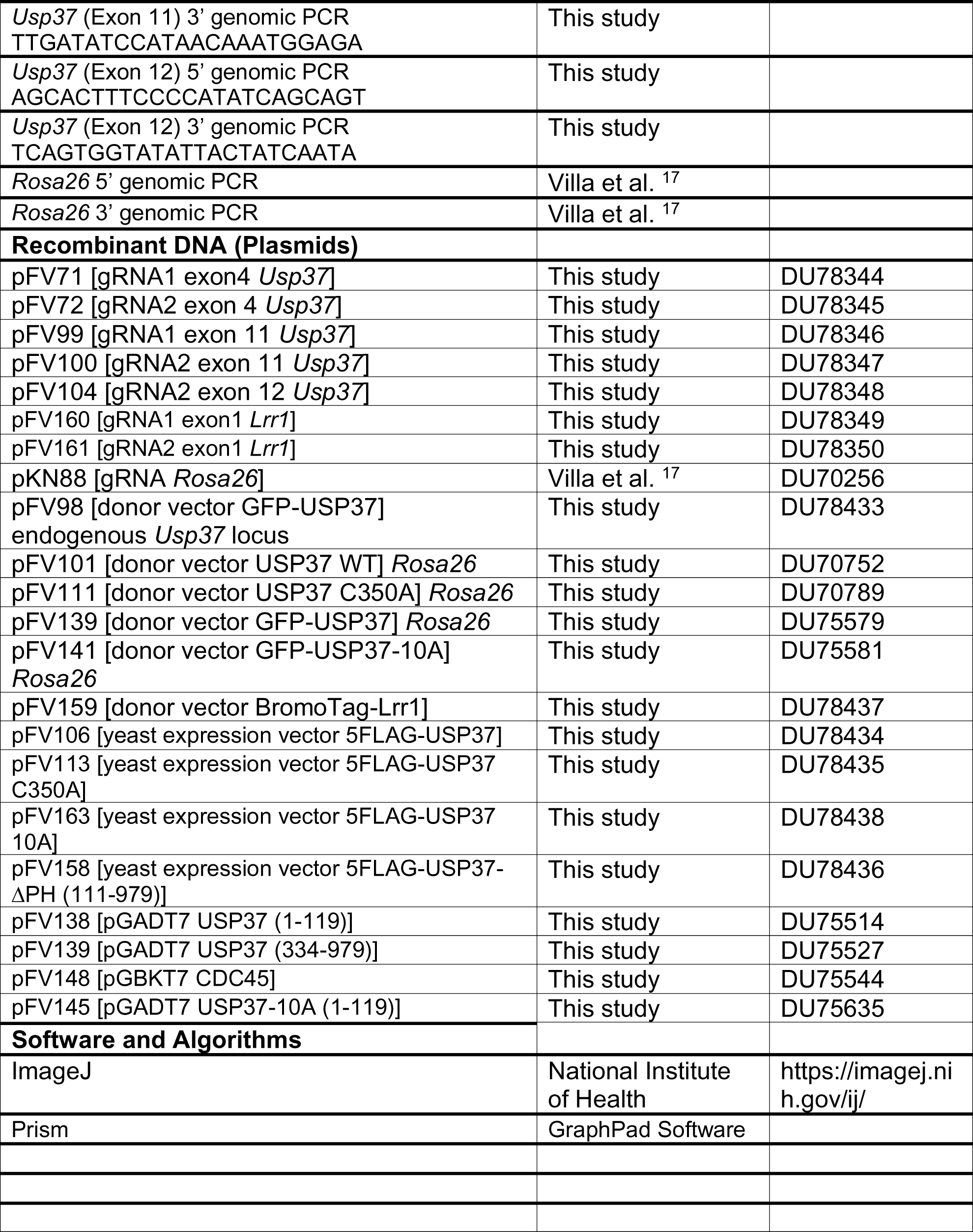
Reagents and resources used in this study.

